# JAMMIT Analysis Defines 2 Semi-Independent Immune Processes Common to 29 Solid Tumors

**DOI:** 10.1101/2021.08.31.458339

**Authors:** Emory Zitello, Michael Vo, Shaoqiu Chen, Scott Bowler, Vedbar Khadka, Thomas Wenska, Peter Hoffmann, Gordon Okimoto, Youping Deng

**Affiliations:** Department of Molecular Biosciences and Bioengineering, University of Hawaii at Manoa; Department of Quantitative Health Sciences, John A. Burns School of Medicine, University of Hawaii at Manoa; SNR Analytics, Inc.; Department of Medicine, Division of Infectious Diseases, Weill Cornell Medicine, New York, NY, USA; Department of Cell and Molecular Biology, University of Hawaii at Manoa

**Keywords:** : Gene signature, immunophenotype, immunotherapy, innate immunity, innate lymphoid cells, monocyte, pan-cancer, squamous cell carcinoma, T_H_17, Warburg

## Abstract

Immunophenotype of solid tumors has relevance to cancer immunotherapy, as not all patients respond optimally to treatment utilizing monoclonal antibodies. Bioinformatic studies have failed to clearly identify tumor immunophenotype in a way that encompasses a wide variety of tumor types and highlights fundamental differences among them, complicating prediction of patient clinical response. The novel JAMMIT algorithm was used to analyze mRNA data for 33 cancer types in The Cancer Genome Atlas (TCGA). We found that B cells and T cells constitute the principal source of variation in most patient cohorts, and that virtually all solid malignancies formed three hierarchical clustering patterns with similar molecular features. The second main source of variability in transcriptomic studies we attribute to monocytes. We identified the three tumor types as T_C_1-mediated, T_C_17-mediated and non-immunogenic immunophenotypes and used a 3-gene signature to approximate infiltration by agranulocytes. Methods of *in silico* validation such as pathway analysis, Cibersort and published data from treated cohorts were used to substantiate these findings. Monocytic infiltrate is found to be related to patient survival according to immunophenotype, important differences in some solid tumors are identified and deficiencies of common bioinformatic approaches relevant to diagnosis are detailed by this work.

## Introduction

Interest in defining the mechanisms responsible for immunotherapeutic response versus resistance have spawned numerous studies. Solid tumors have a fairly similar clinical response rate to immunotherapeutics, and recent results have suggested that T cell polarization may be a critical element in determining patient response to different pro-inflammatory monoclonal agents^1^.

Early findings suggested that Type I immune responses (those involving high levels of interferon-γ, interleukin-2 (*IL-2*), perforins and granzymes) are generally necessary for favorable response to anti-cancer monoclonal-based immunotherapy. Interaction of programmed death-1 (PD-1) with its cognate ligand, PD-L1, inhibits *IL-2* production and inhibits expansion of T_H_1-polarized T cells, while blockade of PD-1 and other inhibitory ligands can result in expansion of anti-tumor T cell clones^2^. High numbers of regulatory T (T_reg_) cells can compete for *IL-2* and thus dampen an inflammatory milieu while supporting the polarization of T_H_17 cells^3^ and promoting survival of CD8^+^ T_C_17 cytotoxic effector T cells. Tumoricidal activity within the tumor microenvironment is mediated by release of perforins and granzymes upon T cell recognition of antigens displayed on major histocompatibility molecules expressed by host tissues and by natural killer cells, which recognize lipid antigens. The activity of innate immune cells has been less extensively studied in cancer immunophenotype and pro-inflammatory therapies. However, a number of investigations have implied that “M2” macrophages, “tumor-associated macrophages,” and “myeloid-derived suppressor cells” can be deleterious to patient health^4—8^.

The Cancer Genome Atlas^9^ (TCGA) repository was launched to address the serious nature and high prevalence of the disease with a standardized, well-planned and multi-modal inquiry among large cohorts of diagnosed patients. The project was made possible in part through advances in high-throughput technologies and data acquisition, and public access to this data has prompted analyses from various approaches. An expanding number of these have been published, pertaining to both individual cancer types^10, 11^ and to pan-cancer studies^12, 13^.

Small cohort studies have investigated cancer immunophenotype with promising results^14, 15^. However, few analyses have characterized the immunologic similarities among solid tumors that account for the observed broad spectrum of activity of monoclonal anti-cancer agents across multiple tumor types, and no known bioinformatic method has clearly defined the role of CD4^+^ T_H_17 and CD8^+^ T_C_17 T cells in cancer from transcriptomic data of whole tissue. Also, the role of the cells of the innate immune system has seldom been included in analyses of tumor immunophenotype, impeding development of a broad consensus with regard to the role of tumor-associated macrophages (TAM’s) and “M1” and “M2” macrophages in cancer. Members of our group previously developed JAMMIT (Joint Analysis of Multiple Matrices by Iteration^16^) and applied the technique to both experimentally-acquired data and the publicly-available TCGA database. Here we show results based on JAMMIT analysis of multiple tumor types from TCGA. We relate this to previous work with collaborators that was collected during an experimental study of biliary tumors which utilized a fluorocholine PET tracer^17^ and profiled global gene expression concordantly with microarray technology. Utilizing these unique approaches to stratify patient cohorts, we provide evidence of the basis for response to immunotherapy for which standard bioinformatic approaches have often sought but have not detailed a strong consensus, while also suggesting new directions in diagnosis, therapy and inquiry.

## Results

### Loadings Plots of JAMMIT Signatures Define a Characteristic “3-Peaks” Signature Common to Solid Tumors

We used JAMMIT to analyze messenger RNA (mRNA) data available in TCGA (https://www.cancer.gov/tcga). JAMMIT uses a “sparse” version of the singular value decomposition to reduce a pre-processed data matrix to a small signature of several hundred genes with high variation among samples by means of a sparse matrix approximation of rank-1. Although JAMMIT is highly adaptable for performance of multi-omics studies, analyses in this publication were primarily unimodal.

Because JAMMIT produces a rank-1 approximation, some discretion was used in manual adjustment of clustering schemes resulting from hierarchical clustering of samples according to JAMMIT solutions.

Upon plotting JAMMIT solutions, a highly conspicuous “3-peaks signature” emerged. Organization within this 3-peaks signature reflects arrangement of gene symbols into sorted alphabetical order. Thus, a small peak of CD (cluster of differentiation) markers, chemokines and chemokine ligands^18^ (with abbreviations beginning with CCL, CCR, CXC and the like) comprise a peak at the left of the loadings plot (***Figure 1A-F***), while genes of VDJ recombination^19^ form two distinct peaks located near the center and at the far right of the loadings plots, respectively. The central peak (corresponding to immunoglobulins) consistently contributed most to the overall solution, while T cell receptor-related genes consistently represented the second most variable element among cohorts. A less conspicuous fourth peak relating to F_C_ receptors^20^ was also visible on most plots. To further clarify the significance of the common gene expression among tumors suggested by the loadings plot, several heatmaps of these genes have been included (***Figure 1— figure supplement 1A-F***). These plots testify to a higher or lower clonal diversity and relative abundance of B cells and T cells among samples independent of tissue type. These findings also demonstrate that the greatest source of variability in individual patient samples measured by bulk mRNA expression in 29 of 31 solid tumor types was related to immune system function and the relative abundance or absence of its cellular components.

**Figure 1.**
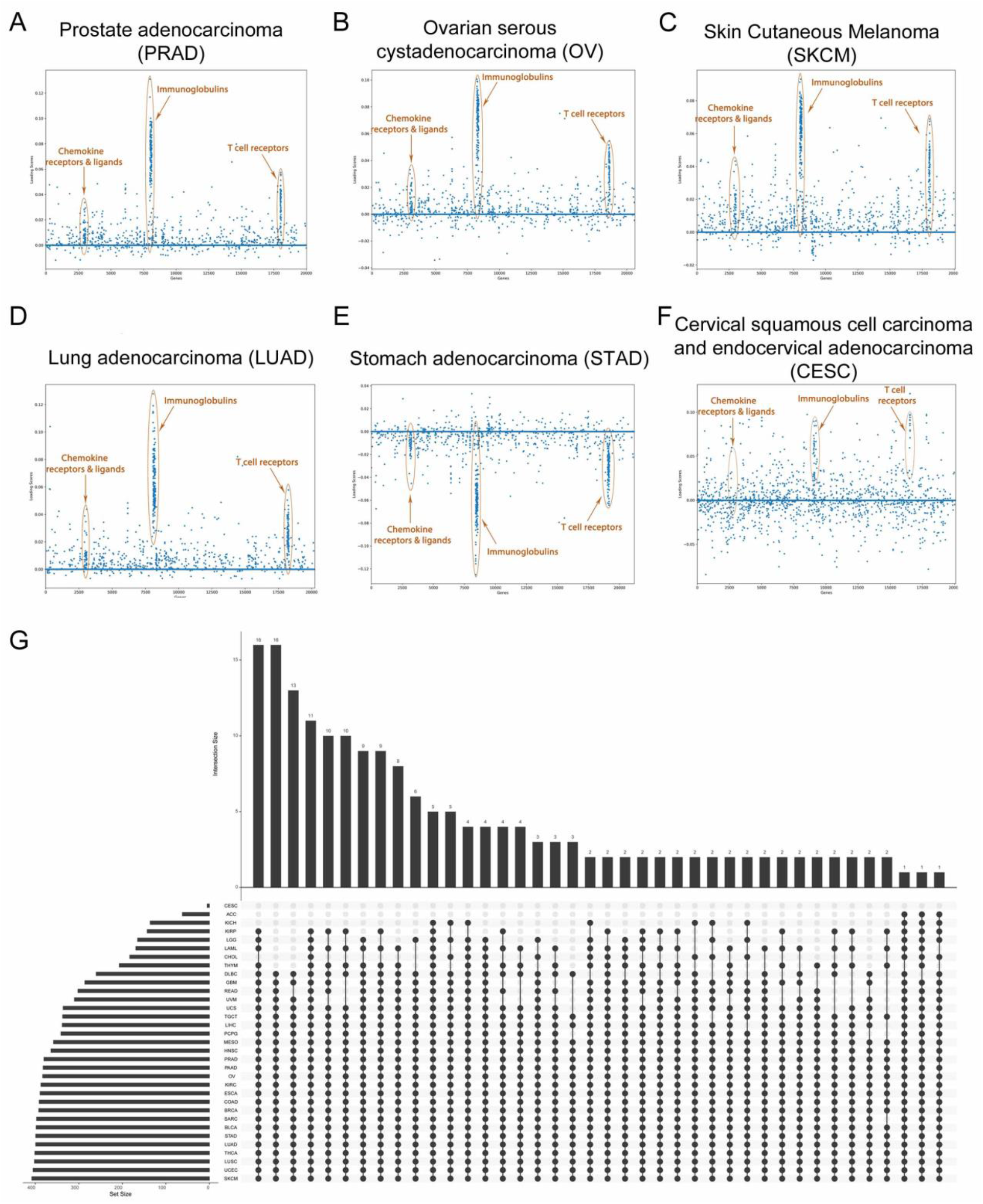
Similarity of Molecular Features of Solid Tumors as Determined by JAMMIT Solution Signatures. (A) through (D) When gene symbols are sorted alphabetically, loadings plots for JAMMIT solutions show similar features for nearly all tumor types in The Cancer Genome Atlas (TCGA). (E) The loadings plot for stomach adenocarcinoma and several other cancers were inverted, but a high degree of overlap among the genes was nonetheless observed. (F) The JAMMIT solution for cervical cancer was somewhat atypical, with the “3-peaks” signature embedded in a noisier background. (G) The UpSet figure shows intersection of unimodal JAMMIT gene signature solutions for 33 cancers in TCGA. Only genes found in at least 25 JAMMIT solutions were included in producing this figure. Exclusion of the genes involved in the process of VDJ recombination in B cells and T cells and lowering the threshold to genes found in at least 15 cancers produced another figure (not shown) with a much smaller intersection size. **Figure supplement 1.** Relationship of JAMMIT Solution-defined Genes to Hierarchical Cluster.

**Figure supplement 1.**
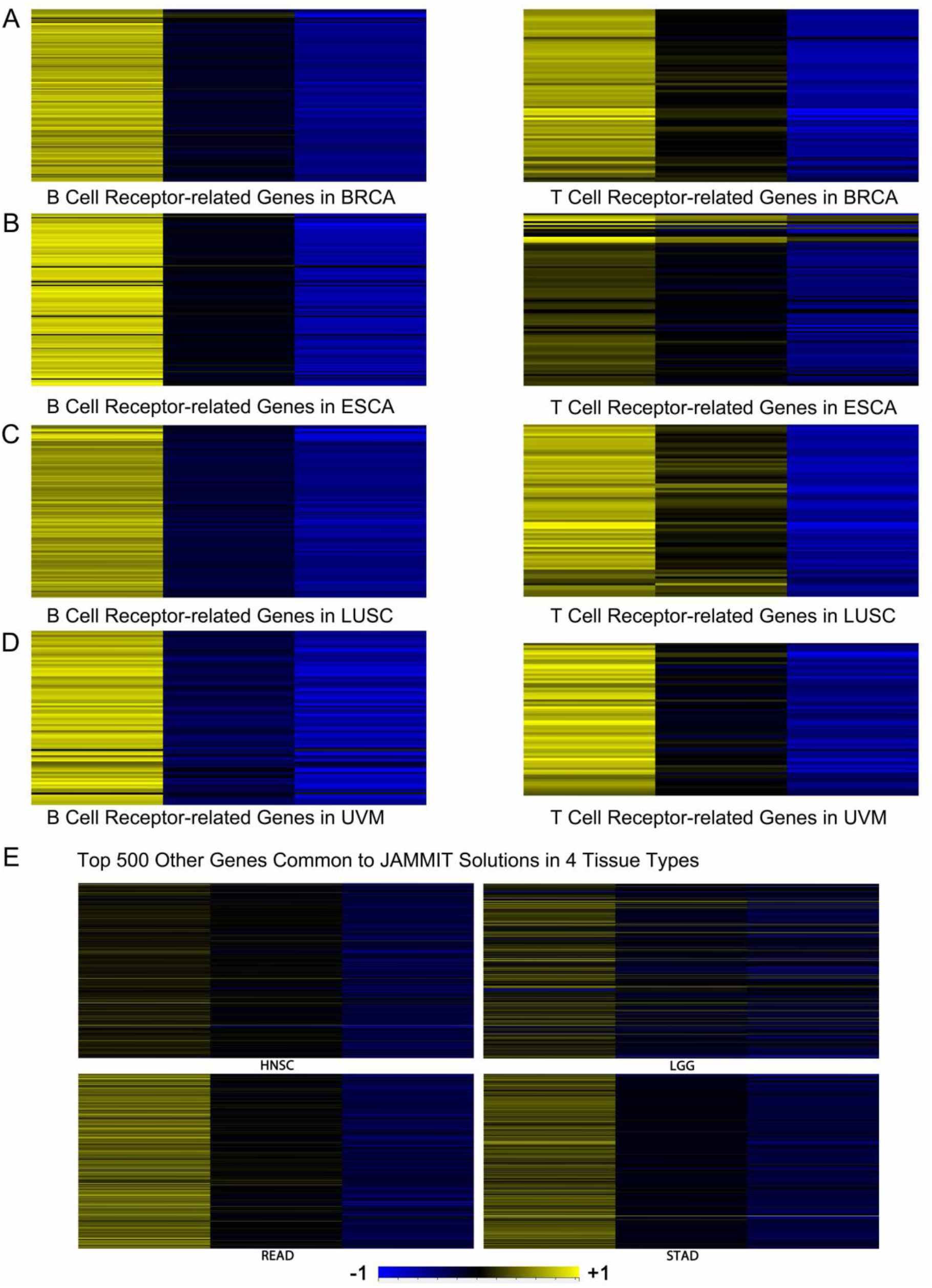
Relationship of JAMMIT Solution-defined Genes to Hierarchical Cluster. **A** to **D** Average gene expression levels of the genes of VDJ recombination in 4 TCGA cohorts. **A** BRCA, **B** ESCA, **C** LUSC, **D** UVM. **E** Per-gene normalized data representing the top 500 genes common to JAMMIT solutions *excluding* the genes of VDJ recombination in 4 TCGA cohorts. HNSC, READ, LGG and STAD are depicted.

Individual JAMMIT solutions had high overlap and with noted exceptions, were primarily immunologic in character. Another subset of genes identified in typical JAMMIT solutions represented mRNA expressed by tissue at the anatomic site of tumor occurrence. This formed the basis for a dichotomy between T_C_1-mediated tumor microenvironments and non-immunogenic tumors. To emphasize common features of these solutions, we produced an UpSet figure^21^ that shows their intersection. As suggested in ***Figure 1G***, solutions showed a modest intersection of genes across a range of tumor types. Many genes involved in the larger intersections shown are related to VDJ recombination; several immunotherapeutic targets and cluster of differentiation markers are also included. To make the character of JAMMIT signatures more coherent, we isolated genes common to the 33 JAMMIT solutions. After removing genes of VDJ recombination, average expression values of the top 500 genes common to JAMMIT solutions were displayed on heatmaps (see also ***Figure 1 –supplementary file 1***). Ingenuity Pathway analysis (IPA) of these 500 genes characterized them as associated primarily with T_H_1 and T_H_2 activation pathways. These heatmaps show that JAMMIT isolates genes that separate cancers into distinct phenotypes independent of anatomical location.

### Functional Analysis of JAMMIT Signatures

To identify biological processes captured in the JAMMIT signatures, we again used IPA. Though precise details varied, IPA returned generally similar results for all cancers. Three pathways significantly involved in all 33 cancer types suggested universal presence of neutrophils and monocytes. In most cohorts, IPA also identified the T_H_1, T_H_2 and T_H_17 activation pathways as top canonical pathways with *p* < 0.05. These results are presented as a bar chart (***Figure 2A***) and also as an UpSet figure (***Figure 2— figure supplement 1A***). Several cohorts stood as relative outliers, but a majority of tumor types intersected in some 30 to 40 molecular pathways.

**Figure 2.**
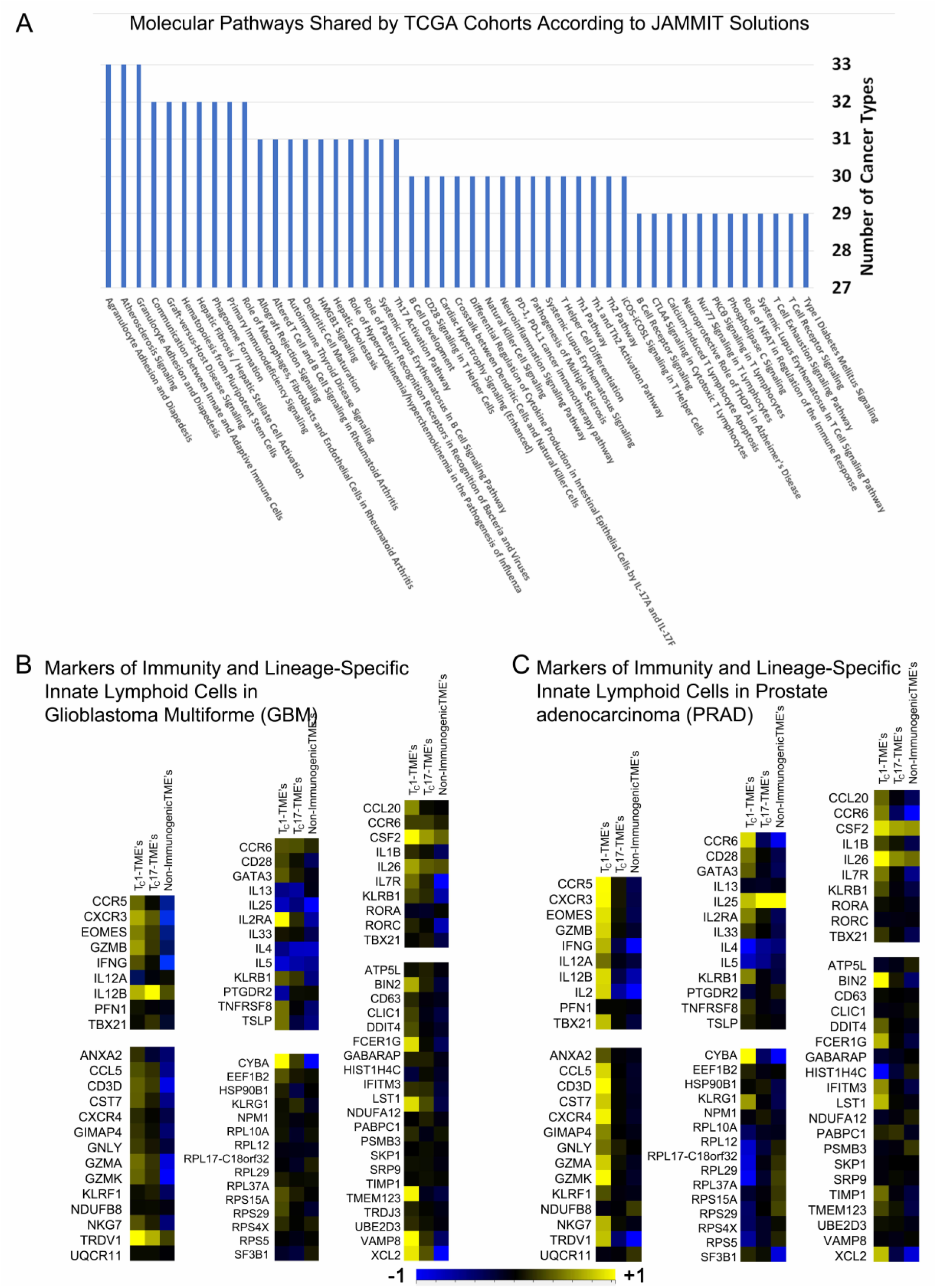
JAMMIT solutions isolate similar molecular features shared by different cancers. (A) Ingenuity Pathway Analysis identified significant cellular and molecular pathways associated with loadings derived from JAMMIT solutions for TCGA mRNA databases. All pathways shown had significance of at least *p* < 0.05. Note that the overwhelming predominance of immune-related pathways indicates similar immune processes are common features of tumors irrespective of cancer or tissue type. (B and C) Features of Type I, Type II and Type III Immunity in JAMMIT solution-derived hierarchical clustering schemes. In each pair of heatmap panels, general immune markers are shown at the top, while lineage-specific markers of innate lymphoid cells are shown below. Differential expression occurs primarily among markers of Type I immunity and secondarily among markers of Type III immunity, but high expression is restricted on average to tumors mediated by Type I immunity. **Figure supplement 1.** Functional Analysis of JAMMIT Signatures for the Various Cancer Types.

**Figure supplement 1.**
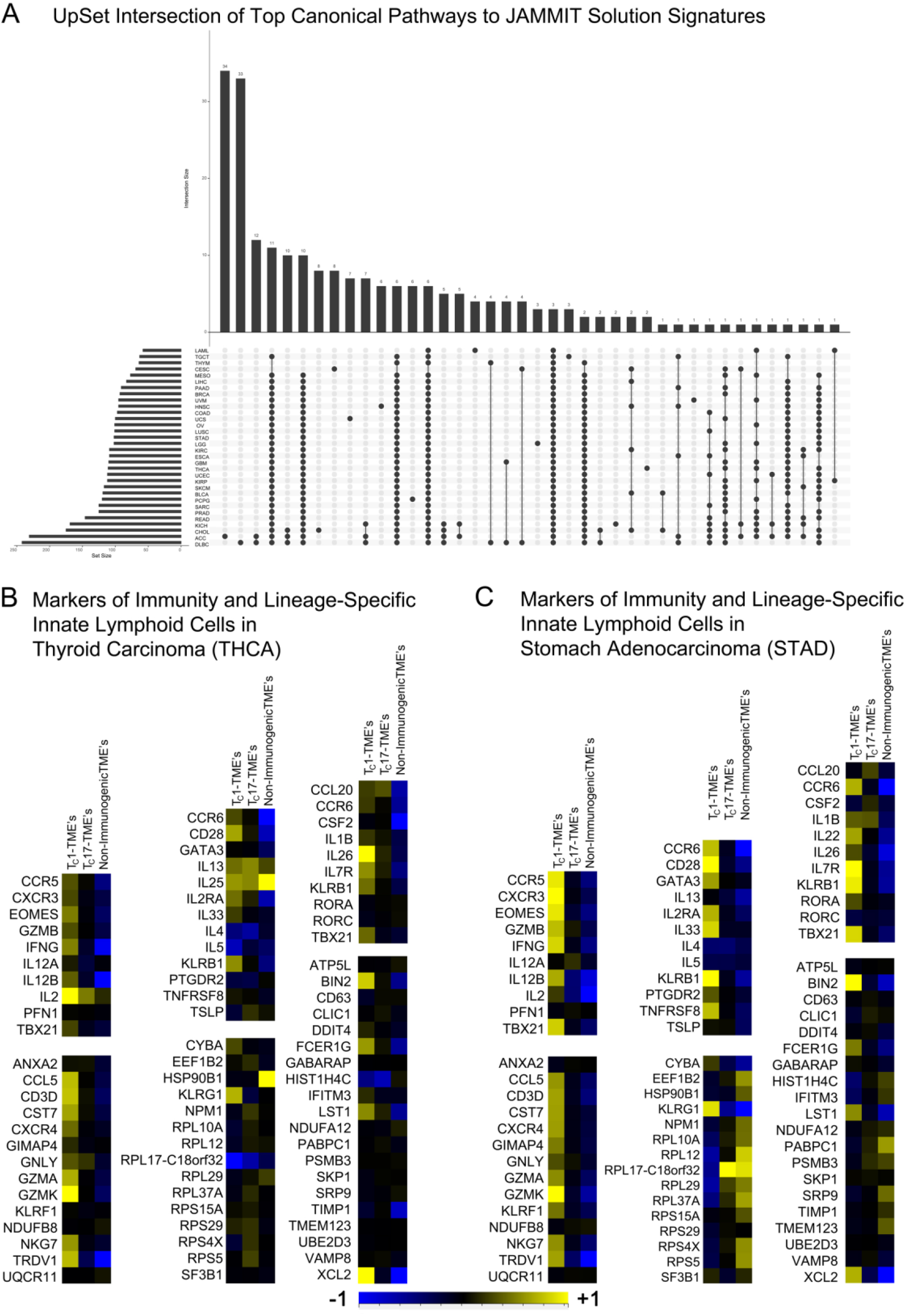
JAMMIT solutions isolate similar molecular features shared by different cancers. A The UpSet intersection of all IPA-identified top canonical pathways derived from JAMMIT solutions for TCGA mRNA databases. All pathways included had a threshold of significance of *p* < 0.05. B and C Features of Type I, Type II and Type III Immunity in JAMMIT solution-derived hierarchical clustering schemes for C THCA and D STAD. In each pair of heatmap panels, general immune markers are shown at the top, while lineage-specific markers of innate lymphoid cells are shown below. Note the similarity to Figures 2B and 2C.

To validate these findings, we used signatures of genes expressed by innate lymphoid cells^22^ and markers of Type I, Type II and Type III immune responses^23^. As shown in ***Figure 2B,C*** and ***Figure 2— figure supplement 1B,C***, these results support IPA’s identification of one cluster of samples as mediated by Type I immunity. However, genes of Type II and Type III immunity did not show consistent, high differential expression of similar quality. We did not interpret this finding as suggesting that Type II or Type III immunity are strictly irrelevant to solid tumors. Rather, we interpreted these results to imply that *only Type I immunity can be recognized in standard approaches to bioinformatic inquiries of solid tumors*. Ordinary analysis of differential expression was not sufficient to distinguish all immune responses in the tumor microenvironment. It is important to mention that no known epidemiological studies have drawn strong ties between T_H_2-mediated disease such as allergy or asthma to high incidence of cancer. The presence of CD4^+^ T_H_17 T cells and CD8^+^ T_C_17 cells have been difficult to characterize in bioinformatical analyses of whole-tissue specimens (see for example publications by Thorsson, *et al*^24^, Hoadley, *et al*^13^, or for an analysis more similar to the present work, see Zhou, *et al*^15^). We therefore tentatively identified the clusters present as T_C_1-mediated, T_C_17-mediated and non-immunogenic immunophenotypes, then characterized these clusters further.

### Hierarchical Clustering According to JAMMIT Solutions Defines 3 Phenotypic Clusters for Multiple Solid Tumors

Reduced matrices of mRNA values to genes determined by JAMMIT solutions produced heatmaps with conventional appearances (***Figure 3A-F***). Following after the similarity of JAMMIT solutions and loadings plots, hierarchical clustering revealed similar biological processes in multiple solid tumors. B cell-expressed genes clustered tightly along the rows with high differential expression among clusters. Genes of T cell receptors (including those of γδ T cells^25-^) ordinarily clustered closely with genes expressed by myeloid cells. Inflammatory markers, B cell-related transcripts, and nearly every immune cell-specific transcript, including markers of Type II and Type III immunity demonstrated highest expression in a single cluster.

**Figure 3.**
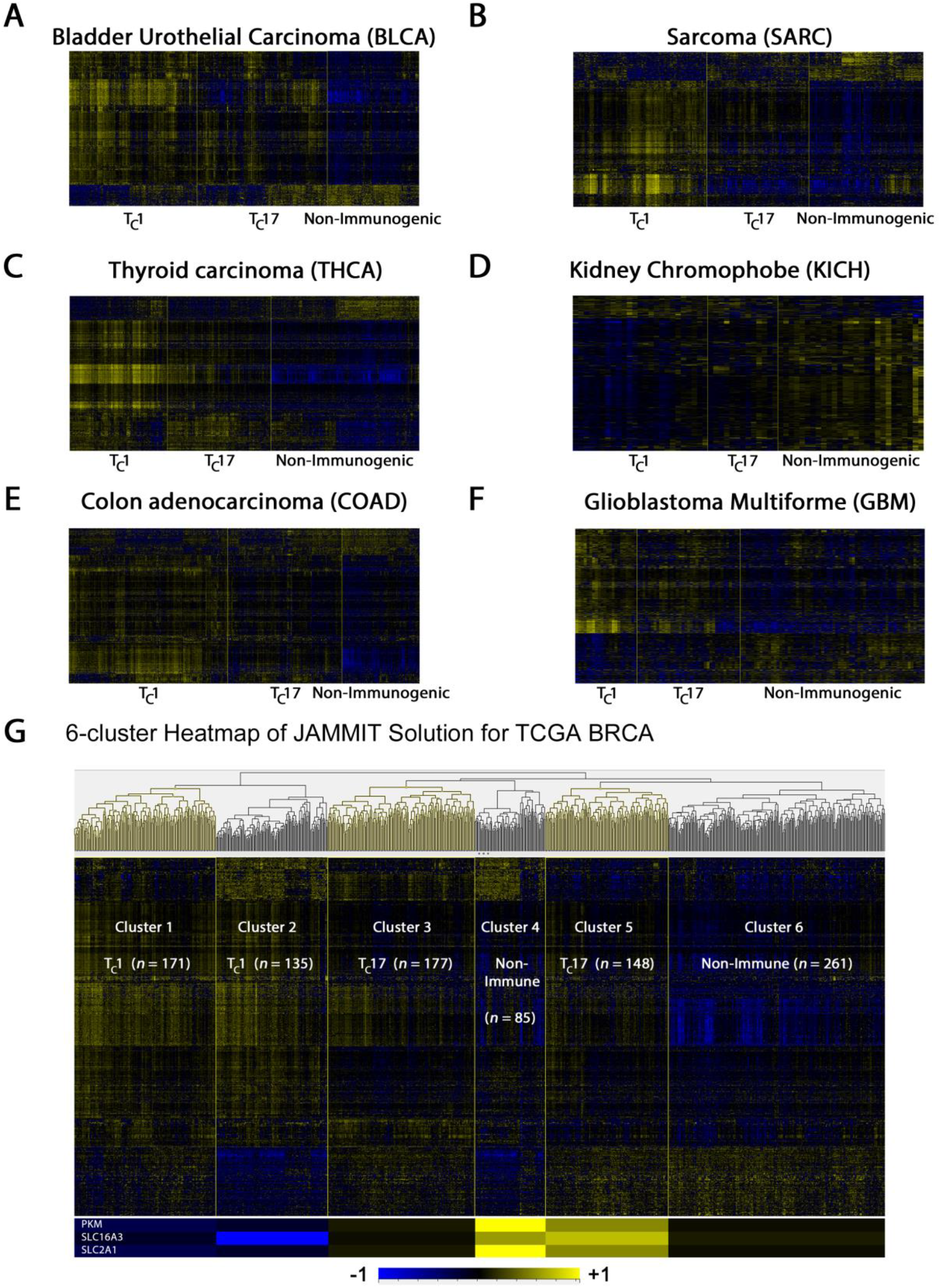
Representative Examples of Clustergrams Produced by Clustering Algorithms of JAMMIT Solutions to Solid Tumors in TCGA. Hierarchical clustering of JAMMIT solution signatures generally resulted in 3 readily distinguished clusters of samples. In each tissue type shown, the clusters with high gene expression of JAMMIT signatures (red) show evidence of relatively dense immune cell infiltrate, while clusters with the lowest expression (green) are consistent with what have been identified as cancers harboring mutations of β-catenin (see text). (A) BLCA (B) SARC (C) THCA (D) KICH (E) COAD (F) OV (G) Hierarchical clustering of the JAMMIT solution for the BRCA cohort produced 6 distinct clusters; the lower heatmap panel shows relative expression levels of a 3-gene “Warburg” signature. **Figure supplement 1.** Agranulocyte Adhesion and Diapedesis in BRCA and Immunophenotype Distribution.

**Figure supplement 1.**
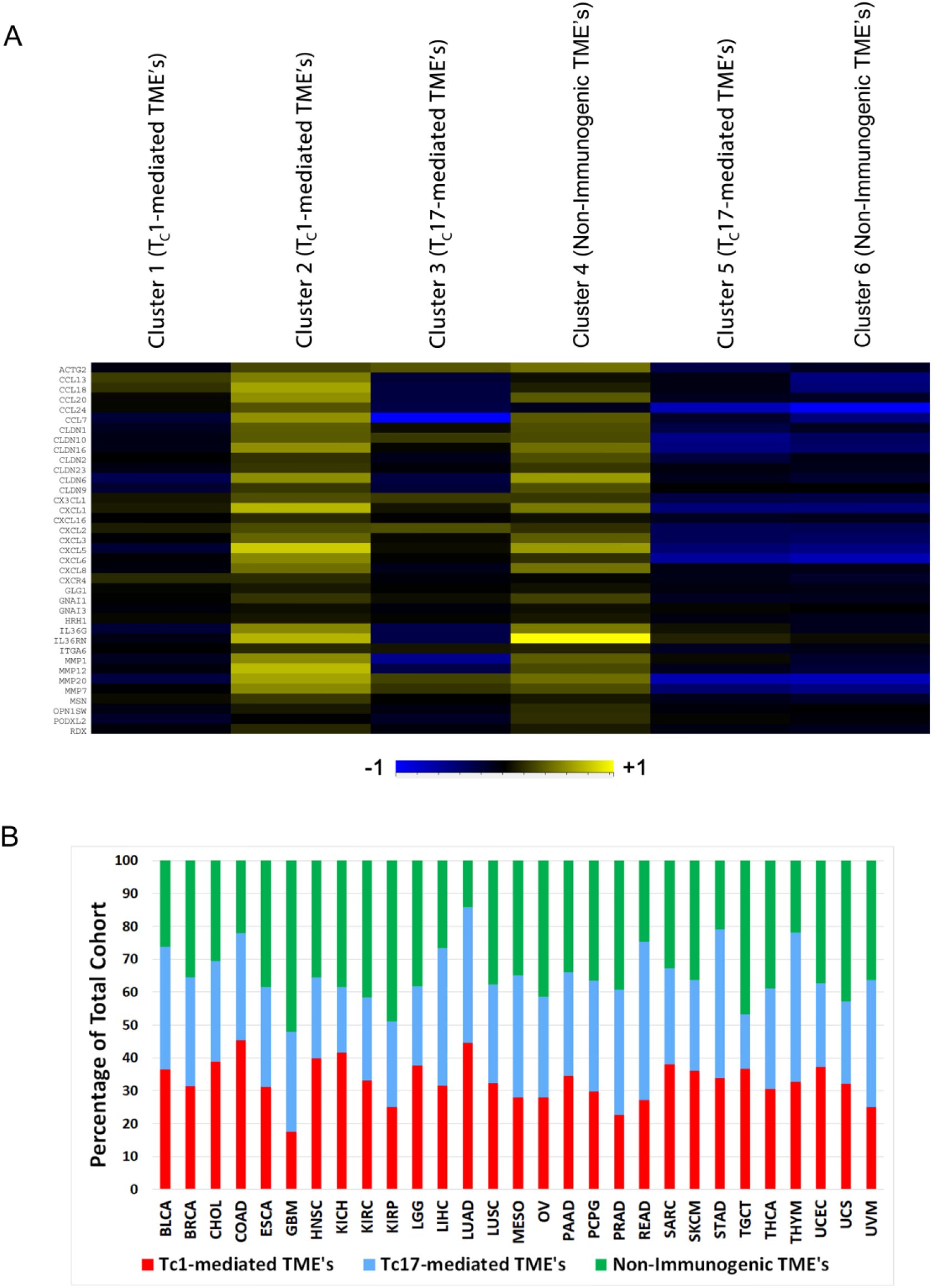
A. Agranulocyte Adhesion and Diapedesis in six subclusters of samples from TCGA BRCA. The heatmap depicts relative mRNA expression of 37 genes of IPA’s canonical pathway related to gene expression of agranulocytes. The clusters are produced by hierarchical clustering of 977 samples according to a 1056-gene JAMMIT signature and show high differential expression primarily among T_C_1-mediated and non-immunogenic samples. **B** The relative proportion of estimated immunophenotype is plotted as a percentage of all samples for each of 29 solid tumor types. In the case of ACC and also for CESC, no division of samples into an analogous clustering scheme was attempted.

Conversely, highest expression of a small number of JAMMIT-defined genes typically characterized non-immunogenic samples. Intermediate expression of most JAMMIT solution-defining genes was generally found in the “middle cluster,” which became a focal point of our studies. In every dataset with the distinctive “3-peaks” pattern, this middle cluster showed modest expression of B cell and T cell transcripts, chemokines and cytokines compared to the inflamed cluster, but relatively high expression of the same genes by comparison with the immunologically silent specimens. Ranking and enumeration of highly-expressed genes in this middle cluster produced results that were neither intuitive nor distinct. Comparison of only a few datasets made it clear that there was no strong or obvious biological process related to high differential expression in these weakly-immunogenic samples. The estimated proportions of the 3 phenotypic clusters in each of 29 solid tumor types for which such a clustering was feasible is shown in ***Figure 3— figure supplement 1B***.

A subtle feature of the hierarchical clustering patterns is highlighted for 977 cases of breast cancer (***Figure 3G***). Standard bioinformatic procedures produced three large clusters of samples with distinct molecular features. However, a closer view reveals that there are actually six hierarchical clusters present, with immunologically similar clusters interrupted by juxtaposition of mathematically similar clusters. As observed at the lower portion of the clustergram, the second and fourth clusters share a similar feature in relatively low expression of a number of genes at the bottom of the heatmap, but are distinguished from one another by a higher clonal diversity and abundance of B cells in the second cluster. The fourth cluster of tumors has molecular features similar to the cluster at the far right and separates two immunologically related clusters on either side. Samples within clusters 3 and 5 differ in the expression of genes related to the “Warburg effect,” as do the first two clusters at the left of the heatmap. These differences proved to be related to the degree of granulocyte and agranulocyte infiltration according to IPA (***Figure 3— Supplementary file 1***).

This leads to discussion of another finding of our work performed with PET imaging.

### A 3-gene Warburg Signature is Correlated with Monocytic Infiltrate, Classical Markers of “M2” Macrophages and Survival

Positron emission tomography has been applied in clinical imaging for over 25 years, and has often utilized a fluorodeoxyglucose tracer^26^. With collaborators^17^, we employed an experimental fluorocholine tracer and found that some tumors had high uptake of glucose but poor avidity for fluorocholine. We identified these as “Warburg” tumors, after Nobel laureate Otto Warburg. Warburg recognized that abnormal fermentation was present, and claimed that cancer was characterized by upregulated glucose uptake and aerobic glycolysis^27^. In more modern studies, this Warburg effect has been characterized in terms of enzymes involved in glycolytic pathways^28, 29^. This has also been more recently called the “Reverse Warburg Effect^30, 31^,” in recognition of the involvement of multiple cell types in establishing a “lactate shuttle” among cancer-associated fibroblasts^32^ and malignant tumor cells. Through application of JAMMIT to microarray data supervised by a PET scan enabled pharmacokinetic parameter, feature reduction techniques allowed derivation of a 3-gene signature (*PKM*, *SLC16A3* and *SLC2A1*) that preferentially identified tumors with poor uptake of fluorocholine.

We sought to explore this signature’s relevance. Upon examination of TCGA data ordered by ranked expression of this signature, we found that samples with high and low expression differed significantly in expression of classical markers of “M2” macrophages^33, 34^ across cohorts (***Figure 4A,B***). Cibersort deconvolution of mRNA data validated this finding with estimates of increased monocyte infiltration (***Figure 4C-F***). In concord with findings of others who have explored the relevance of myeloid-derived suppressor cells^6^ and “M2” macrophages^7^, we found that this 3-gene signature had prognostic value in multiple patient cohorts (see representative examples in ***Figure 4G,H***).

**Figure 4.**
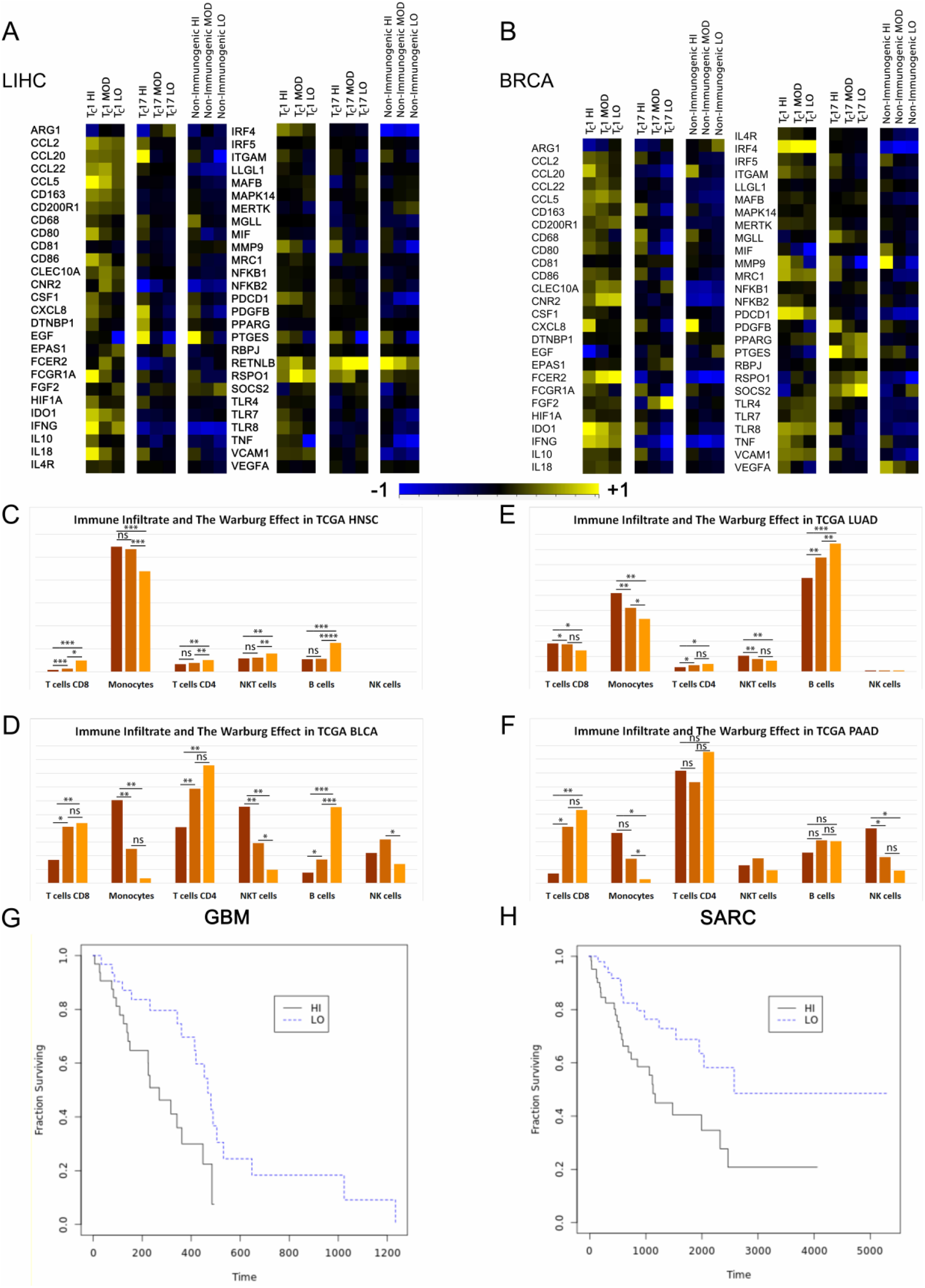
Correlates of the Warburg Effect. (**A** and **B**) Relative gene expression levels of some popular markers of M1, M2 and Tumor-associated Macrophages as stratified by JAMMIT solutions and ordering according to a “Warburg Index” defined by a 3-gene signature. (**A**) Per-gene normalized heatmaps of some popular markers of M1 and M2 macrophages and TAM’s as seen in immunophenotypic clusters of 260 samples from TCGA LIHC database. The samples have also been subdivided according to high, moderate and low Warburg Index, as indicated. (**B**) Similar representation of data as seen in 977 cases of breast cancer in TCGA that have been hierarchically clustered according to a 1056-gene JAMMIT signature. (**C** to **F**) Cibersort estimates of immune cell distribution at high, moderate and low levels of the Warburg effect. (**C**) HNSC, (**D**) BLCA, (**E**) LUAD, (**F**) PAAD. (**G** and **H**) Kaplan-Meier survival analysis for the GBM and SARC cohorts. (**G**) GBM : χ2 = 7.76, *p* = 0.0053, (**H**) SARC χ2 = 7.96, *p* = 0.0048. **Figure supplement 1.** Markers of Activation and Differentiation and JAMMIT Solutions. **Figure supplement 2.** Immunophenotype, the Warburg effect and patient survival.

**Figure supplement 1.**
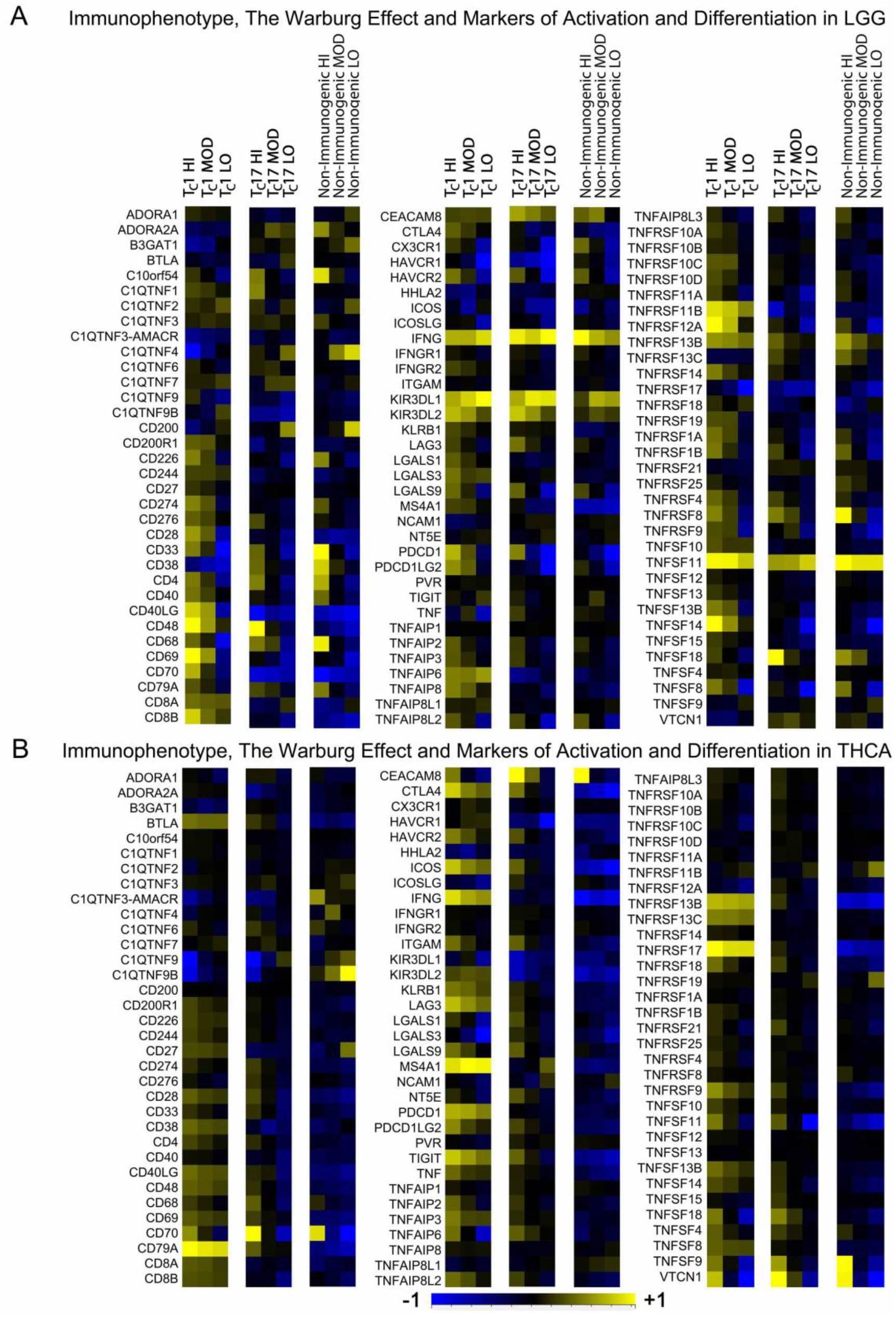
Relative Gene Expression Levels of Markers of Activation and Differentiation Other Markers of Immunity as Stratified by JAMMIT Solutions and Ordering According to a “Warburg Index” Defined by a 3-gene Signature. **A** Genes corresponding to immune cell surface markers, popular and novel immunotherapeutic targets, and their cognate ligands as expressed in 507 cases of low grade glioma from TCGA. Samples were hierarchically clustered by a JAMMIT signature and ranked as high, moderate or low by Warburg Index. **B** A similar panel of genes as expressed in 501 cases of thyroid carcinoma from TCGA. Both heatmaps were produced using per-gene normalization.

**Figure supplement 2.**
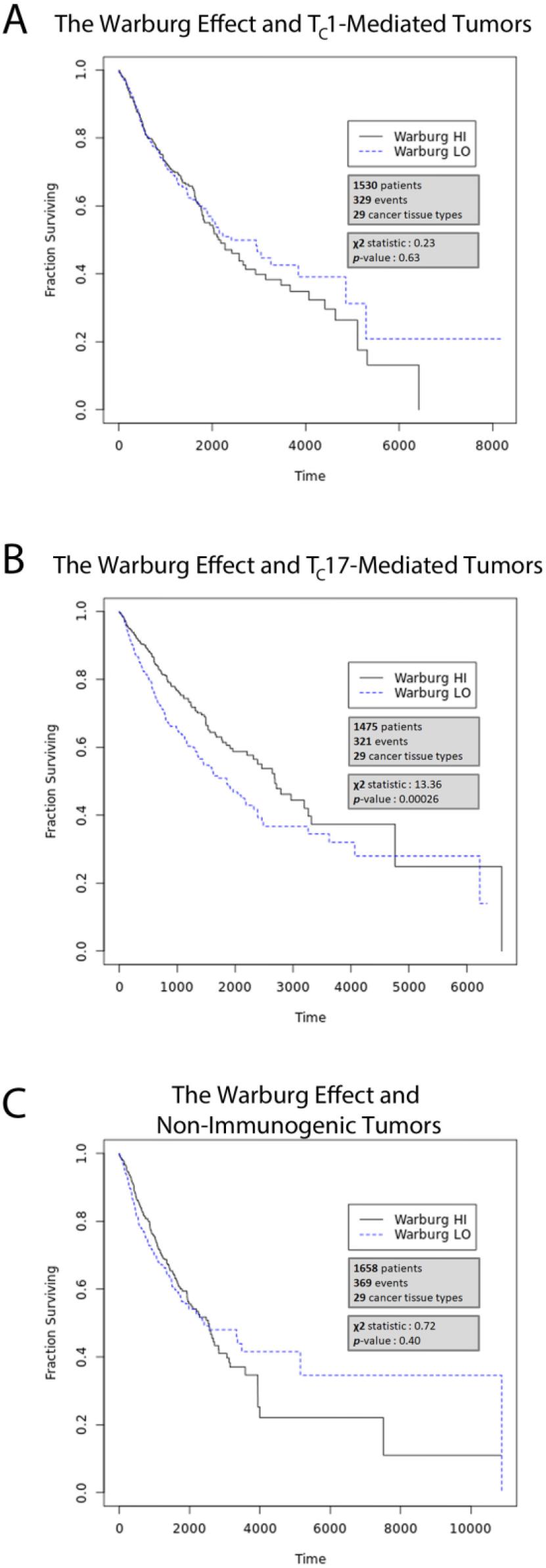
The Warburg effect and its relationship to immunophenotype and survival. The relationship of the Warburg effect and survival for patients with solid tumors and similar immunophenotype are shown. The number of patients of each immunophenotype are indicated. All analyses were performed using Mantel-Haenszel variant of the log-rank test. Statistics for individual cohorts are as follows: (**A**) T_C_1 HI vs. LO, χ2 = 0.23, *p* = 0.63; (**B**) T_C_17 HI vs. LO, χ2 = 13.36, *p* = 0.00026; (**C**) Non-Immunogenic HI *vs.* LO, χ2 = 0.72, *p* = 0.40.

Some have suggested that variation in myeloid cells subsets may be associated with different patient outcomes^35, 36^. From our vantage point, studies of the tumor microenvironment only provide information concerning the relative abundance or scarcity of myeloid cells. According to JAMMIT solutions, differences in immunophenotype overshadowed any differences in innate immune cell infiltrate; thus we emphasize once again that the primary difference in global transcriptome among patients with solid tumors in most cohorts is defined by immunophenotype, while the differences in innate immunity are a cause of *secondary* variation. However, we did note that when viewed in isolation from other immunophenotypes, patients bearing T_C_17-mediated tumors who had high levels of monocytes within the tumor microenvironment had an unexpected survival benefit compared to those who had few (***Figure 4—figure supplement 2***). Whether this reflects a functional difference in the myeloid cells in these patients is unclear. Speculatively, the prognostic difference may reflect a difference in the cytokine influence of Type III immunity upon innate immune cells or immune cell-cell interactions. Our interpretation of a lack of Type II immunity in cancer does lead us to favor the opinion that eosinophils are not prominent in ordinary solid tumors, however.

### The Warburg Effect and Markers of Activation and Differentiation

Observed modulation of macrophage gene expression by lactate has been reported^37^. Similarly, lactate can affect gene expression of other immune cell subsets^38^. These facts and others led us to inspect a wider panel of standard and novel immunotherapeutic targets for modulation by this Warburg effect in other cohorts. Similar to markers of M1 and M2 macrophages and TAM’s, transcript expression of these genes and other well-known immune cell surface markers mirrored the expression of B cell and T cell receptor gene distribution with regard to the immunophenotypic clusters, but exhibited more modest differential expression (***Figure 4 – figure supplement 1A,B***). Given that many of the targets of modern immunotherapy are expressed by T cells, we found it particularly noteworthy that high levels of the Warburg effect accompanied high expression of many of these ligands. This was often true despite a modest reduction in the number of infiltrating T cells (see next section and ***Figure 4C-F***).

### The T_C_1-Mediated Tumor Immunophenotype

In the case of T_C_1-mediated TME’s, strong evidence provided by IPA analysis suggested that the cluster was characterized by Type 1 immunity much as defined by Annunziato, *et al*^23^. High expression of genes specific to Type 1 innate lymphoid cells in multiple tissue types (see again ***Figure 2C*** and section on Innate Lymphoid Cells below) generally corroborated this finding. At the molecular level, these clusters showed high differential expression of hallmarks of Type 1 immunity, such as *IFNG*, *IL-2*, *IL-12*, T-bet, and eomesodermin in most tissue types. In spite of IPA’s identification of “T_H_1-T_H_2 pathway” as a top canonical pathway, markers of Type 2 immunity such as *CRTH2*^39^, *IL-4*, *IL-5*, *IL-13* and GATA3^40^ often showed no consistent, organized pattern of differential expression in these clusters. Highly conspicuous was the finding that essentially all genes related to B cells (*CD79A*, *MS4A1*, *MZB1*, *PAX5*, *etc.*) and B cell-chemotactic factor *CXCL13* exhibited highest average expression in this cluster. Similarly, the T_C_1-mediated cluster contained the highest average expression of more than 50 T cell transcripts, consistent with highly-infiltrating T_C_1 CD8^+^ and T_H_1 CD4^+^ T cells that display activity against intracellular pathogens^41^. Though a better understanding of the relationship between T_reg_ cells and T_H_17 CD4^+^ T cells has emerged^42^, the finding of *FOXP3* as a marker of T_reg_ cells having their highest abundance within these clusters is novel. Most of these findings applied uniformly across all solid tumors. These findings indicated a generally higher level of immune cells in the TME with a predominant signature characterized by T_C_1 cells.

### The Non-Immunogenic Tumor Immunophenotype

For the third cluster of samples with a “cold” appearance on heatmaps, we investigated mutations of β-catenin based on previously published evidence^43, 44^ an established high prevalence of β-catenin mutations among multiple tumor types^24^, and molecular features consistent with immuno-exclusive TME’s, *a.k.a.* “immune deserts.” These immune-exclusive microenvironments were present in every tissue type (see again ***Figure 3A-G***) and comprised one-third of the >9,000+ tumors of the 29 solid tumor types studied for which 3 distinct clusters could be identified. Generally, T_C_17-mediated tumors were distinguishable from non-immunogenic counterparts by more than 100 differentially-expressed genes of VDJ recombination. In pairwise comparisons, at least as many of these genes withstood Bonferroni correction in every dataset with at least 200 samples. One exception was found for the low-grade glioma (LGG) cohort, which had a peculiar aberration of low *CXCL13* expression within the T_C_1-mediated cluster. Mutations of genes in the Wnt-β-catenin signaling pathway are known to disrupt cell-surface expression of proteins necessary for normal physiologic T cell adherence to the exterior of malignant cells^43^, which may explain why genes of T cell receptors would be absent from these tumors.

The status of this “non-immunogenic” cluster first became clear upon review of the literature of pertaining to bioinformatical analysis of hepatocellular tumors by both the TCGA research network authors^11^ and very similar results produced by another team^45^. In both cases, the rate of β-catenin mutations among large cohorts of patients with hepatocellular carcinoma was found to be ∼27%, matching rather precisely with hierarchical clustering according to our own JAMMIT-defined solution for hepatocellular carcinoma (LIHC).

Because we did not use raw DNA sequence data in our analysis, we felt it appropriate to further confirm the non-immunogenic nature of these samples with the data available at hand. Based on expression of an 89-gene signature recognized by IPA as pertinent to Wnt-β-catenin signaling, we produced heatmaps of differentially expressed genes for several tumor tissue types. A typical result showing generally higher expression of most genes of the canonical Wnt-β-catenin signaling pathway within T_C_1- and T_C_17-mediated clusters is shown in ***Figure 5 – figure supplement 1A***.

**Figure 5.**
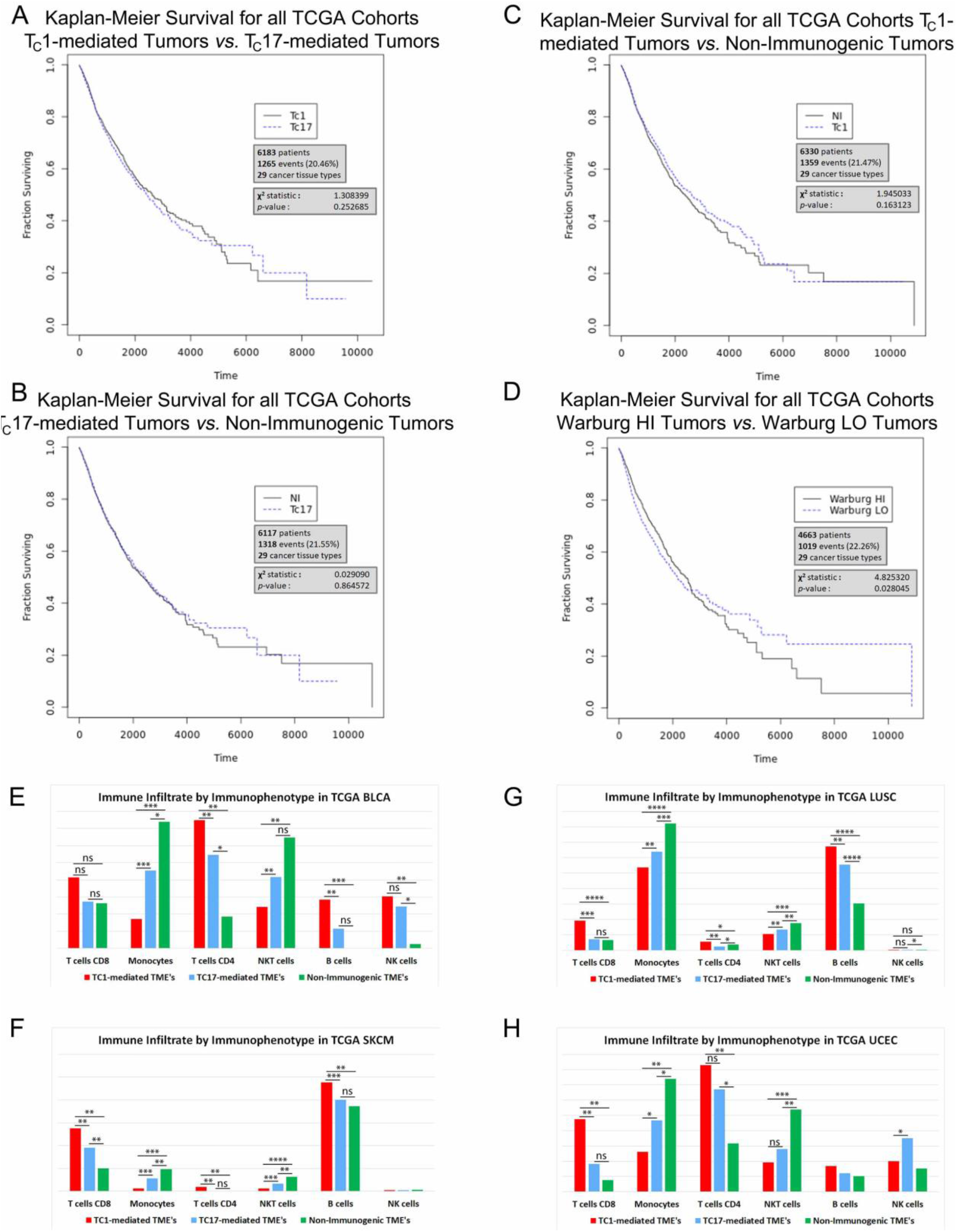
Correlates of Tumor Immunophenotype. The relationship of immunophenotype, the Warburg effect and survival for patients with solid tumors in TCGA datasests are shown. The tumor types and number of patients in each cohort are indicated. Note that only the Warburg effect had prognostic significance in this analysis. All analyses were performed using Mantel-Haenszel variant of the log-rank test. Statistics for individual cohorts are as follows: (**A**) T_C_1 vs. T_C_17, χ2 = 1.31, *p* = 0.25; (**B**) T_C_17 vs. Non-Immunogenic, χ2 = 0.029, *p* = 0.86; (**C**) T_C_1 vs. Non-Immunogenic, χ2 = 1.94, *p* = 0.16; (**D**) Warburg HI vs. Warburg LO, χ2 = 4.82, *p* = 0.028. (**E** through **H**) Cibersort estimates of immune cell infiltrate in representative TCGA datasets. All datasets shown here have been clustered according to JAMMIT solutions with some minor modifications, as alluded to in the text. Note a consistent higher estimate of the number of T cells in T_C_1-mediated tumor samples (red) in nearly every comparison. \**p* < 0.05, \*\**p* < 0.01, \*\*\**p* < 10-5, \*\*\*\**p* < 10-10 . (**E**) BLCA, (**F**) SKCM, (**G**) LUSC, (**H**) UCEC.

**Figure supplement 1.**
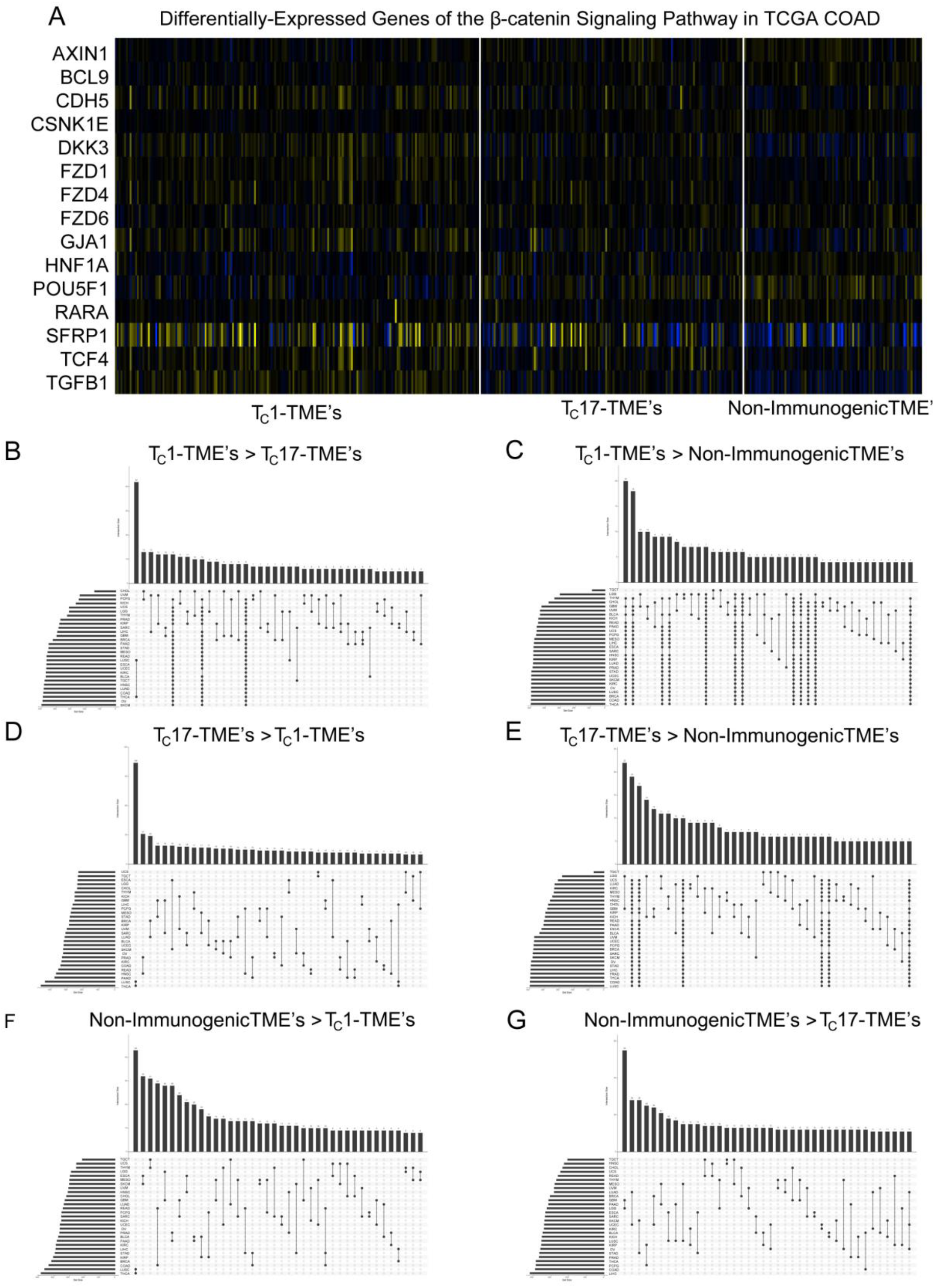
Immunophenotype Characterization. (**A)** 15 differentially expressed genes of an 87-gene pathway signature are shown. All 15 of these genes showed statistically significant differential expression with *p* < 0.05 (using a standard Student’s t-test) in each of 3 pair-wise comparisons among T_C_1-mediated, T_C_17-mediated and non-immunogenic tumors as hierarchically arranged according to a 926-gene JAMMIT signature. Note the generally higher expression of all genes in the two immunogenic clusters and the contrast with non-immunogenic tumors, which suggests a loss of normal function. Of the 87 genes in IPA’s β–catenin pathway, only *SFRP1* was a JAMMIT solution signature-defined gene. (**B** through **G)** Shown in each frame is an UpSet figure produced from pair-wise comparisons of mRNA expression levels of the individual genes within immunophenotypic clusters resulting from hierarchical clustering of JAMMIT solutions. The top 500 differentially-expressed genes for each cluster-wide comparison contributed to a signature for each dataset. Note the similarity of the intersections between (**B)** and (**E)**, which reinforces the notion that many of the same genes of statistically significant differential expression in T_C_1 *versus* T_C_17 comparisons are of similarly high expression in T_C_17-mediated tumors compared to non-immunogenic tumors; thus a “3-layer” stratification exists.

### The T_C_17-Mediated Tumor Immunophenotype

The molecular similarities shared by T_C_1-mediated tumors included a long list of immune cell transcripts demonstrating high expression confined to that cluster, but the middle cluster was not as simply identifiable as being infiltrated by T_C_17 and T_H_17 T cells. IPA analysis failed to identify the T_H_17 pathway among the most highly significant top canonical pathways associated with this cluster in multiple tissue types. T_C_17/ T_H_17 signature genes (*CD73*^46^, *DPP4*^47^, *RORA*, *RORC*^48–50^) and genes related to TH17 function (*IL-6*^51, 52^) showed little or no differential expression across most tumor types. Another well-known marker of T_H_17 CD4^+^ T cells, *IL-17*, exhibited little differential expression in TCGA cohorts and has been shown to have neutrophils as its primary physiological source of relevance^53^.

However, at the bulk level, these tumors were clearly distinguished on heatmaps (***Figure 3A-F***). The differences already noted in the distribution of genes of VDJ recombination pointed to an unequivocal and consistent difference in the abundance of adaptive immune cells attributable to a phenotypic distinction. Recalling that genes of high differential expression within the weakly immunogenic clusters of multiple cohorts showed little to no overlap, we proceeded to analyze these samples with the assumption that they differed from both T_C_1-mediated samples and non-immunogenic tumors. After clustering each of the datasets according to its corresponding JAMMIT solution, with small manual adjustments to hierarchical clustering results, we characterized the middle cluster for each cancer type in a series of six pairwise comparisons. Average transcript expression level and a Student’s t-test applied to the entire transcriptome allowed construction of 6 UpSet figures (***Figure 5 – figure supplement 1B-G***). We chose to use an arbitrary limit of 500 genes for each pairwise comparison to construct these figures. These data (represented in ***Figure 5 – figure supplement 1B, E***) show that T_C_17-mediated tumors have a gene expression profile which is similar to T_C_1-mediated tumors when compared to non-immunogenic tumors, but reflects the gene expression profile of a non-immunogenic tumor by comparison with the highly-infiltrated T_C_1-mediated tumors (***Figure 5 – figure supplement 1D, F***).

Explanation of why T_C_17-mediated tumors would have moderate levels of immune infiltrate in whole-tissue specimens came from a study of hepatocellular tumors by Dong-Ming, *et al*^54^. This study showed that IL-17-secreting CD8^+^ T cells were found concentrated at the invading edge of tumors, rather than throughout malignant tissue. This is consistent with the biological function of T_C_17 T cells, which act by inducing cells of host tissues to secrete defensins and other anti-microbial peptides^55^ and enhance proliferation through *MAPK* signaling^56^. In normal physiology the secretion of antimicrobial peptides has an adaptive role, but activation of *MAPK* in cancer is capable of promoting tumor expansion. In the context of Type III immunity, cellular proliferation is further reinforced through IL22 signaling from Type 3 innate lymphoid cells and T_H_22’s^57^ via induction of STAT3 signaling^58^. Thus, low expression of eomesodermin^41^, granzymes and perforins characteristic of T_C_17 cells cause these tumors to be relatively depleted of immune infiltrate while tumor growth is facilitated. Cibersort^59^ estimates validated the lesser abundance of both CD4^+^ and CD8^+^ T cells in T_C_17-mediated tumors in nearly every tissue type examined, except in renal clear cell carcinoma, thyroid and ovarian carcinoma cohorts, where T_C_17 cells or T_H_17 cells were estimated to outnumber T_C_1-polarized cells in one of these compartments (see ***Figure 5E-H***).

Though it was not a primary aim of this study, it is worth noting that immunophenotypic cluster conferred no significant survival benefit in the handful of datasets we examined individually. Again, at the level of the entire TCGA cohort of patient samples, we found that the Warburg effect was prognostic (***Figure 5A-D***).

### Cross-validation with Data from a Treated Cohort

Given the recent findings in pre-clinical models that suggest tumors infiltrated primarily by T_H_17-polarized cells do not respond well to single-agent immunotherapy^1^, we sought to validate our interpretation of the biological significance of the hierarchical clustering we observed in multiple tumors types. To carry out this validation, we downloaded RNA sequencing data from a cohort of melanoma patients with corresponding information on response to therapy from dbGap^25, 60^. Unfortunately, data acquired from these patient cohorts used a sequencing read depth with insufficient coverage to make our JAMMIT solutions of optimal utility in this setting. Therefore, to investigate whether the middle cluster represented tumors infiltrated by T_C_17 and T_H_17 T cells, we took into account published data in which the gene expression profiles of patients were measured using a Nanostring^61^ platform during courses of anti-PD-1 therapy for treatment of metastatic melanoma. As ***Figure 6*** shows, the gene expression pattern of patients classified as having a more favorable response to therapy in melanoma is more consistent with that of samples identified by JAMMIT analysis as T_C_1-mediated specimens rather than with T_C_17-mediated tumors in several cohorts. This was confirmed using Fisher’s exact test. Though some of the patients in this small cohort were not treatment-naïve, this analysis suggests that Nanostring may be a highly cost-effective platform for measuring gene expression in this context when compared to deep-sequencing of whole tissue specimens.

**Figure 6.**
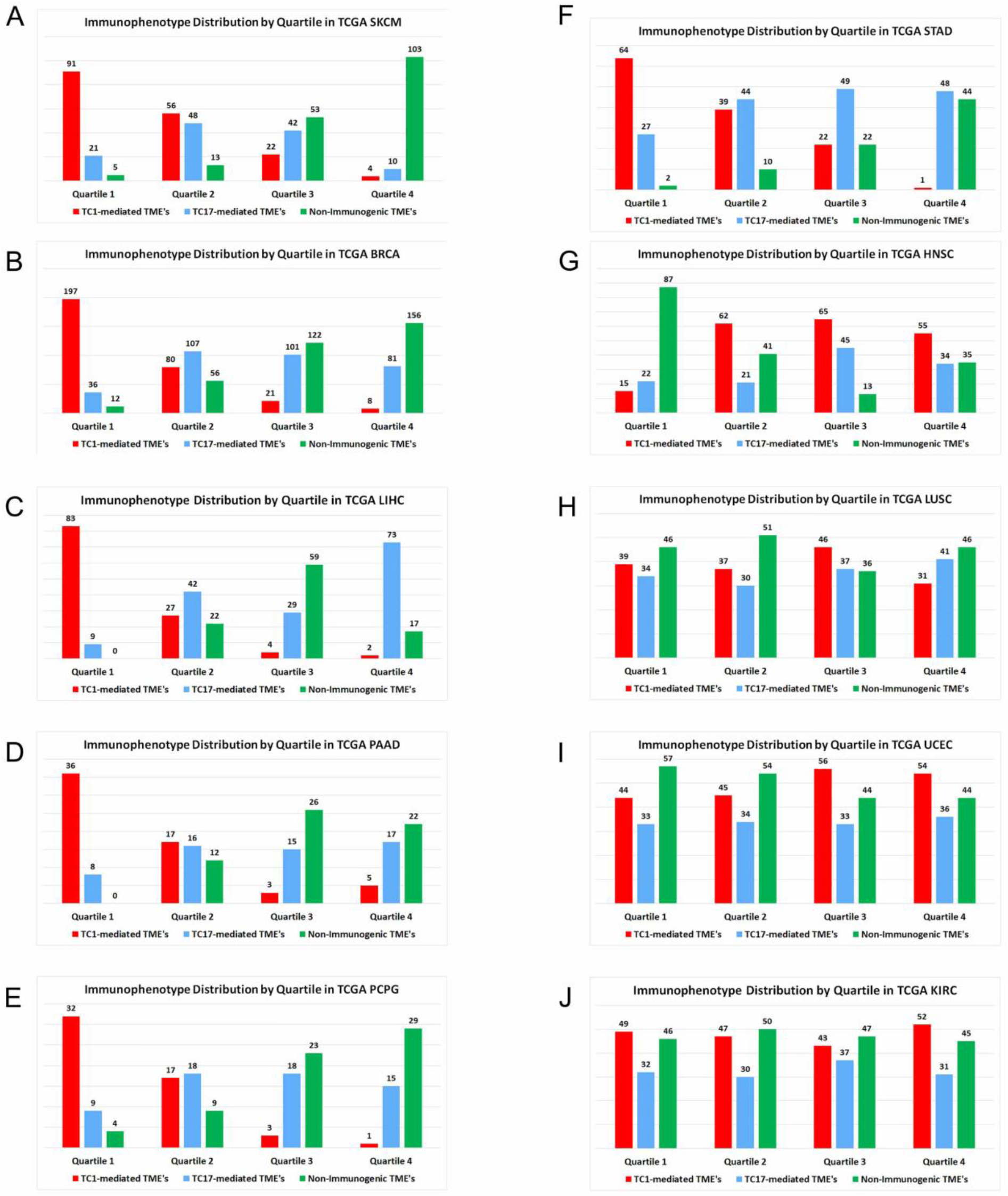
Distribution of patient samples according to rank-order of singular value decomposition scores. The plot shows the per-quartile distribution of patient samples when ranked by expression of a 266-gene signature which distinguished responders to anti-PD-1 immunotherapy in a published report. All samples are categorized by immunophenotype according to hierarchical clustering of JAMMIT solutions for the cancer tissue type shown. The number of samples per quartile per category is shown. The *p*-value shown refers to the results of comparison of the number of T_C_1-mediated *vs.*T_C_17-mediated samples in first and second quartiles according to Fisher’s exact test. (**A**) SKCM (total cohort size *n* = 468), *p* = 0.00011 (**B**) BRCA (total cohort size *n* = 977) , *p* = 0.00000000 (**C**) LIHC (total cohort size *n* = 367) , *p* = 0.00000000 (**D**) PAAD (total cohort size *n* = 177) , *p* = 0.0063 (**E**) PCPG (total cohort size *n* = 178) , *p* = 0.0091 (**F**) STAD (total cohort size *n* = 372) , *p* = 0.0020 (**G** through **J**) Several outliers in validation were also noted. The 266-gene signature that characterized clinical responders to therapy in 28 melanoma patients treated with anti-PD-1 agents did not distinguish T_C_1-mediated tumors from T_C_17-mediated tumors as identified by JAMMIT in these cohorts. (**G**) HNSC (total cohort size *n* = 495), *p* = 0.77 (**H**) LUSC (total cohort size *n* = 474) , *p* = 0.15 (**I**) UCEC (total cohort size *n* = 534) , *p* = 1.00 (**J**) KIRC (total cohort size *n* = 509) , *p* = 1.00

In performing these validations on multiple datasets, we also discovered a limitation of gene expression-based approaches used to predict or measure patient response to immunotherapy. As can be seen for the TCGA cohorts for squamous cell cancers of the lung and head and neck, and also for endometrial cancers, the gene signature that characterized patients responding to anti-PD-1 therapy in melanoma did not extend to all cancer types. In order to investigate the potential reasons for this restriction, we therefore deepened our investigation of these outliers and revisited previous analyses.

### Innate Lymphoid Cells and JAMMIT Analyses

Based upon Cibersort estimates that showed high numbers of monocytes and low numbers of T cells for several datasets, we suspected that innate immunity might be a cause of the different gene expression profiles in our validations pertaining to patient response to immunotherapy and the proposed T_C_1-T_C_17-Non-Immunogenic clustering scheme. With reference again to the previous figures showing that a broad range of immune cell-surface markers, immunotherapeutic targets and TAM markers generally showed high expression in T_C_1-mediated tumors and in concert with high levels of the Warburg effect (***Figure 4A,B*** and ***Figure 4 – figure supplement 1A,B***), it was difficult to explain the opposing trend observed in the head and neck squamous cell carcinoma (HNSC) and lung squamous cell carcinoma (LUSC) cohorts (***Figure 7 – figure supplement 1A,B***).

**Figure 7.**
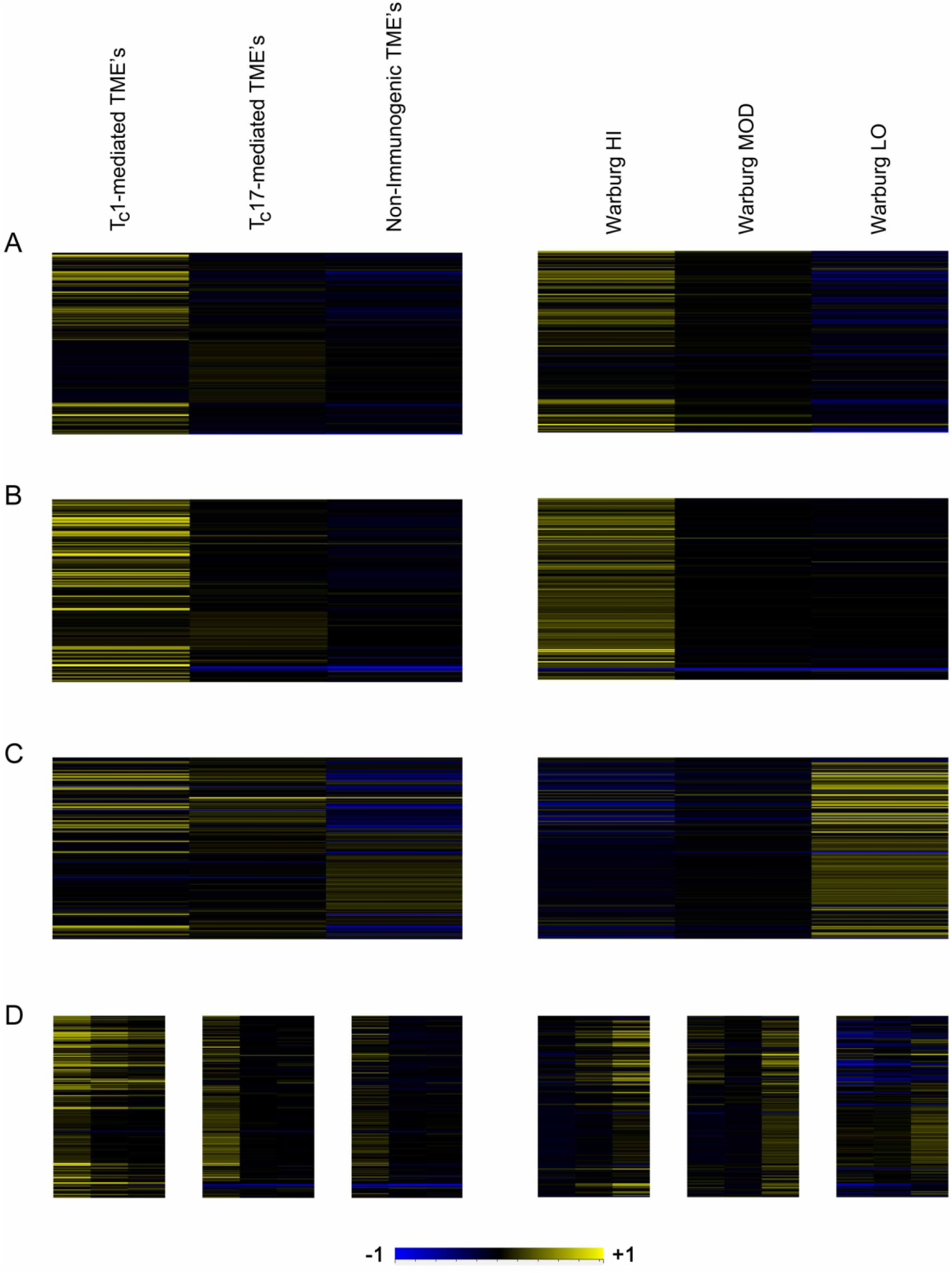
Common innate lymphoid cell signature in solid tumors and in squamous cell carcinoma cohorts. (**A**) through (**C**) Each set of panels depict average expression values of differentially-expressed genes of a 222-gene signature of genes common to innate lymphoid cell lineages clustered either according to immunophenotype (*left*) or Warburg Index (*right*). (**A**) LGG (**B**) LIHC (**C**) HNSC (**D**) Compound immunophenotype (T_C_1, T_C_17 and non-immunogenic panels) and Warburg clustering (high Warburg effect at left, low Warburg effect toward right of each panel) is depicted; at left, LGG exemplifies a typical clustering pattern for non-squamous solid tumors. At right, HNSC shows an opposing trend toward higher number of innate lymphoid cells only at low levels of the Warburg effect. **Figure supplement 1** The Warburg effect and immunophenotype in two squamous cell carcinoma cohorts. **Figure supplement 2** Common innate lymphoid cell signature in solid tumors and in squamous cell carcinoma cohorts.

**Figure supplement 1.**
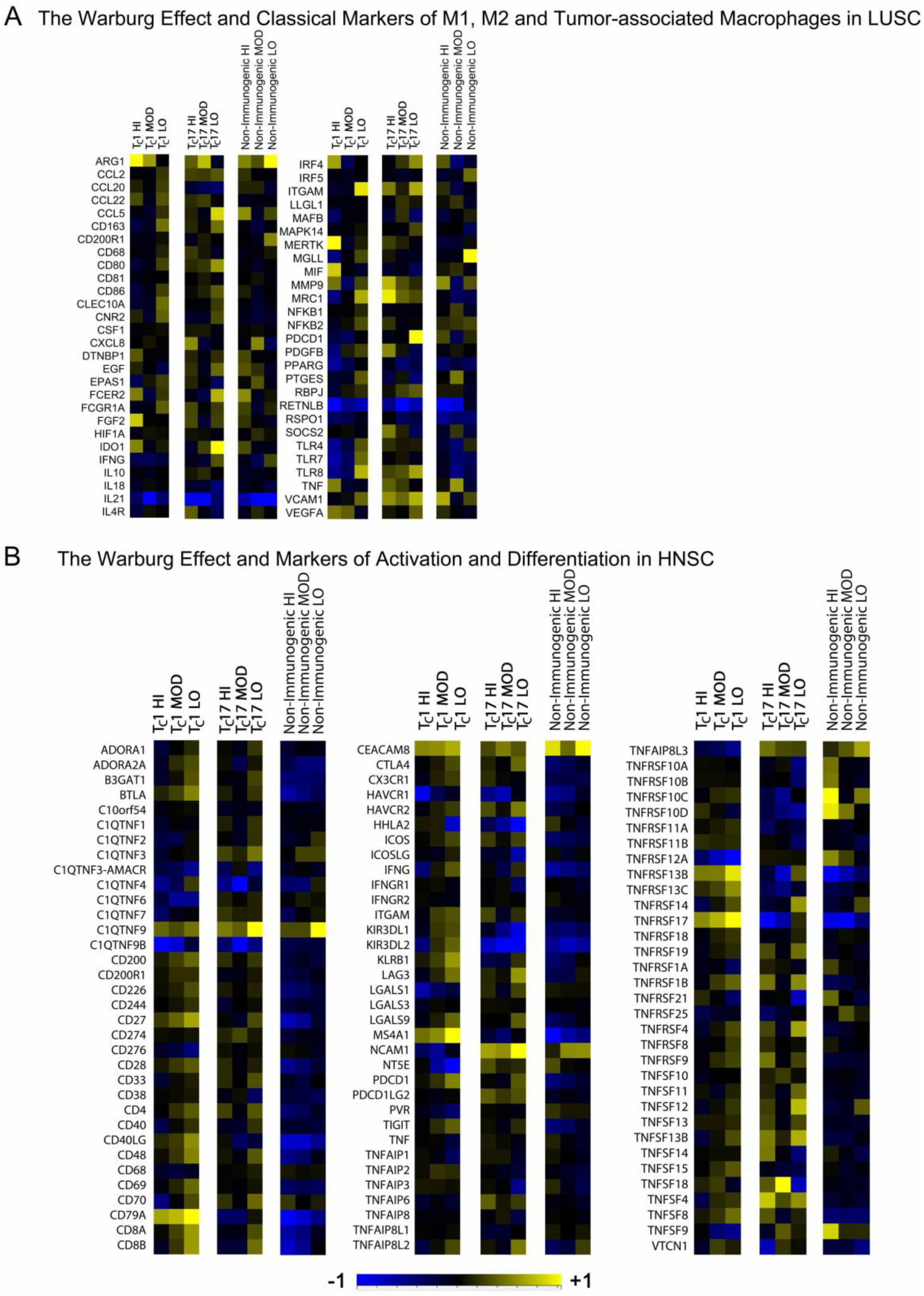
The Warburg effect and immunophenotype in two squamous cell carcinoma cohorts. These cohorts departed from others in demonstrating higher expression of macrophage and B and T cell surface ligands and receptors at low (rather than high) levels of the Warburg effect as determined by the 3-gene signature. (**A**) Popular markers of M1, M2 and tumor-associated macrophages in the LUSC cohort. Compare with Figures 4A and 4B. (**B**) Markers of activation, differentiation and related markers in the HNSC cohort. Compare with Figures S4A and S4B.

**Figure supplement 2.**
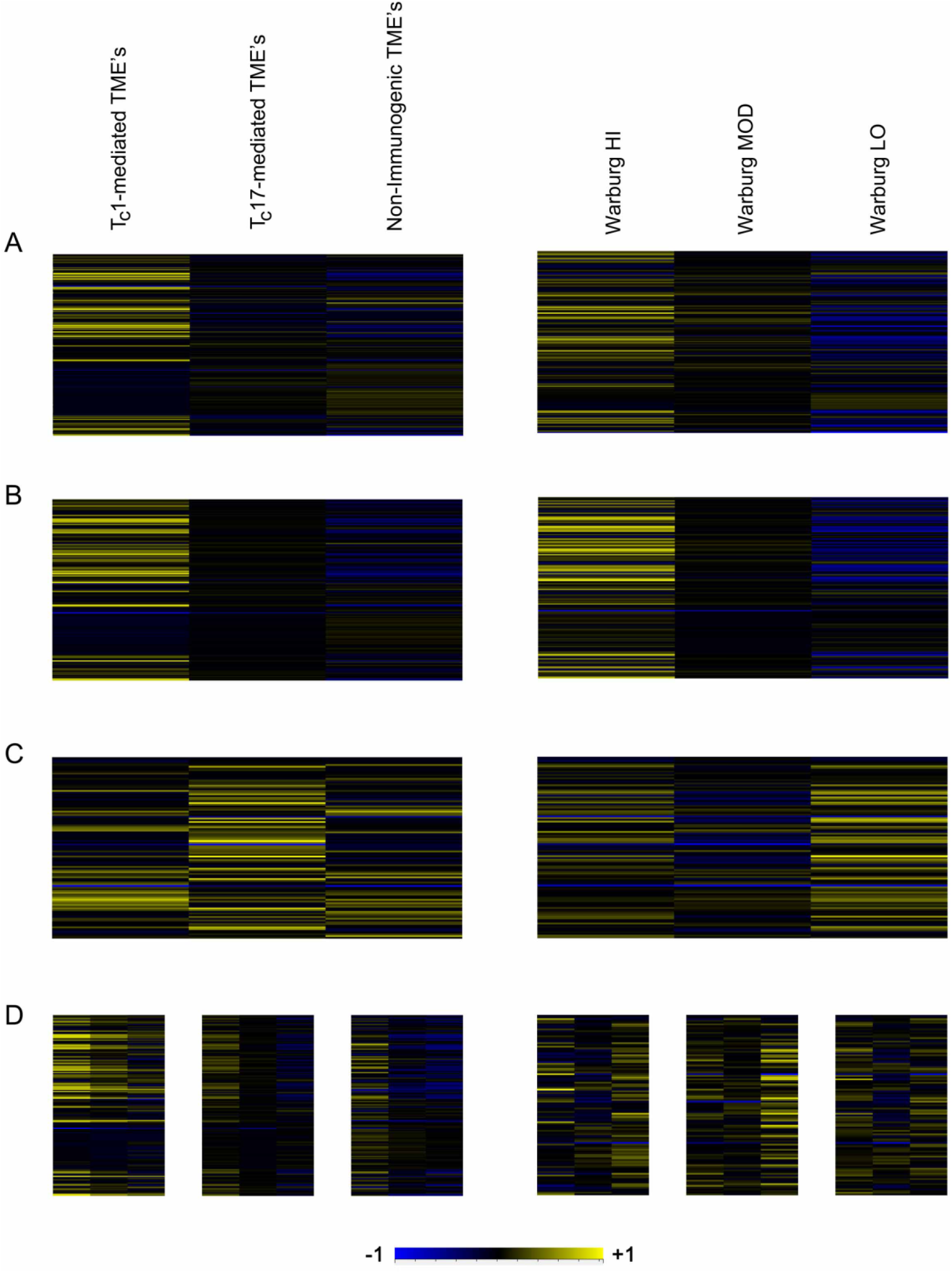
Common innate lymphoid cell signature in solid tumors and in squamous cell carcinoma cohorts. (**A** through **C**) Each set of panels depict average expression values of differentially-expressed genes of a 222-gene signature of genes common to innate lymphoid cell lineages clustered either according to immunophenotype (*left*) or Warburg Index (*right*). (**A**) PCPG (**B**) PRAD (**C**) LUSC (**D**) Compound immunophenotype (T_C_1, T_C_17 and non-immunogenic panels) and Warburg clustering (high Warburg effect at left, low Warburg effect toward right of each panel) is depicted; at left, PRAD exhibits a typical clustering pattern for non-squamous solid tumors. At right, LUSC shows an opposing trend toward higher number of innate lymphoid cells at lower levels of the Warburg effect.

This seemingly unusual departure from other solid tumors was explained by examining expression patterns of genes expressed by innate lymphoid cells. All three major lineages of innate lymphoid cells have been sequenced at the single-cell level^22^, and as genes specific to each lineage were examined (see again ***Figure 2C***), we also took into account a larger signature of genes (222 in all) common to all three lineages. As observed with some large TCGA cohorts, high expression of the genes of this signature again clustered preferentially among samples displaying the T_C_1-dominant immunophenotype (***Figure 7A,B***). Here LGG and LIHC serve as two “prototypical” cohorts (see also ***Figure 7 – figure supplement 2A,B***).

Equivalent analysis of the corresponding cohorts for HNSC and LUSC produced very different results, however (***Figure 7C*** and ***Figure 7 – figure supplement 2C***). From the results of these parallel analyses, we conclude that hierarchical clustering of JAMMIT solutions for TCGA datasets defines a T_C_1-T_C_17-Non-Immunogenic trichotomy of solid tumors for all cohorts of patients with solid tumors exhibiting significant immune activity among at least a certain proportion of samples. However, these immunophenotypes do not always align predictably with the distribution of the cells of innate immunity. The presence of monocytes within tumor microenvironments, particularly in high numbers, appears to have the capacity to disrupt an orchestrated immune response and can skew interpretation of gene signatures.

## Discussion

The search for biomarkers to predict patient response to immunotherapy has often utilized measurements of gene expression taken from the tumor microenvironment. Our studies suggest that one confounding factor in analyses of this type is a failure to take into account the variable number of innate immune cells within the tumor microenvironment being assayed. Another problem in these approaches has been that insufficient read depth has placed bioinformaticians at a great disadvantage in discerning various immune and non-immune phenotypes present in most solid tumor types.

As part of these investigations, JAMMIT analysis allowed us to approximate a T_C_1-T_C_17 dichotomy present in multiple solid tumors from TCGA that previous bioinformatic techniques have not addressed. Tumors with higher numbers of T_H_17 CD4^+^ T cells have been demonstrated in pre-clinical models to be correlated with a lack of effectual treatment response to one or more forms of single-agent immunotherapy utilizing monoclonal antibodies. Thus, the addition of a TGF-β-blocking agent may improve patient response rates by approximately 100% compared to single-agent therapy. In at least one validation dataset outside of the TCGA compendium, JAMMIT was unable to produce such a distinction among the samples of a large patient cohort (*n* = 164). This may be due to insufficient read depth of the mRNA sequencing assays. We also concluded that another reason bioinformatics approaches have failed to identify T_C_17-mediated tumors as a distinct immunophenotype is that they have relied upon high differential expression of genes as a standard to characterize biological processes. However, we found that data produced by our analyses strongly implies the existence of a set of tumors for which gene expression is intermediate *globally* in large cohorts where 3 phenotypes are present in significant numbers.

We also showed that multiple types of cancers of squamous epithelial cell origin likely had high numbers of monocytes, setting them apart from other solid tumor types. Our findings suggest that T cell infiltrate is inherently dampened in these cases, and this is likely to extend to some other cohorts (e.g. squamous cell cancers of the skin). Attempts to validate a gene expression signature proven to predict a response to treatment in melanoma extended well to other tumor types, but not to squamous epithelial tumors. This implies that no universal gene signature exists for responders, suggesting that the basis for patient response to immunotherapy is a characteristic intrinsic to the immune cells themselves (by implication, it is the polarization and receptor sequence of T cells). Previous work with collaborators developed a 3-gene signature in investigation of PET imaging, providing evidence to suggest that this signature has a high correlation with monocytic infiltrate. Thus, other tumor types may potentially be diagnosed through a similar radiographic technique. There is also an implication that infiltration by monocytes can influence the behavior of T cells and may even regulate their numbers. Current literature suggests the concentration of lactic acid in the tumor microenvironment is a probable factor related to this blunting of T cell responses. We also note that the 3 genes in the signature we have used as a proxy for monocyte infiltrate have a close relationship to lactic acid metabolism^,28, 32^, consistent with what has been described as a “Reverse Warburg Effect.” This Reverse Warburg effect has been described in multiple cancers and has been reproduced in murine models^62^.

Finally, we showed by way of this proxy measurement of monocytes that these innate immune cells are likely a primary cause of poor patient prognosis in a cohort of several thousand. This suggests that targeting the innate immune system may be a valuable means of immunotherapy in addition to current methods in use. In particular, reducing bloodborne GM-CSF may represent a feasible approach^63^ to reduce monocyte production by the bone marrow. A caveat to this notion is that some tumor types appear to have very little immune infiltrate other than these monocytes to act as a defense for the host.

## Materials and methods

### Data preprocessing

After download, each individual TCGA cohort was preprocessed for JAMMIT analysis by removal from the expression matrix of all genes with 75% or more entries equal to zero. Data was log-transformed according to a generalized log transformation method^64^ (“glog”), followed by quantile normalization. Centering of each individual matrix row is performed by subtraction of average value from the row.

### JAMMIT analysis

JAMMIT runs for these analyses were set to 1,000 permutations each. Solutions were chosen according to an approximate signature size of 1,000 genes rather than false discovery rate (FDR). This produced an FDR of less than 0.05 for all cohorts except CHOL, THYM, UCS and UVM.

### UpSet plots

UpSet plots were produced using the online tool available as referenced in the text.

### Ingenuity pathway analysis

Top canonical pathways were determined using IPA analysis. Singular value decomposition of matrices formed by JAMMIT solution signatures produced loading values for each gene to be uploaded to IPA for individual cohorts. Only significant pathways with *p* < 0.05 were used in the figures shown.

### Gene signatures

The 3-gene “Warburg” signature was derived during a previous study as referenced in the text. Apart from JAMMIT solution signatures, other gene signatures used in this publication were based on referenced publications, IPA or curated manually.

### Hierarchical clustering

Hierarchical clustering was produced using TreeView with Pearson’s correlation of both rows and columns and complete linkage. Per-gene normalization^65^ of data was employed to create heatmaps of reduced signature size (except for Fig. S1, which is based on average gene expression).

### Cibersort validation

Cibersort validations were conducted with 1,000 permutations using absolute mode and a sample signature matrix file supplied at the Cibersort website among a list of examples.

### Immunotherapeutic efficacy validation

After download of published data, validation of patient response to immunotherapy was performed by segregating responders to anti-PD-1 therapy and non-responders. Differential expression analysis of the panel of genes included in this analysis (including just under 800 genes of primarily immunologic expression) was conducting using a one-tailed homoscedastic Student’s t-test. Transcriptomic mRNA data for significantly differentially expressed genes were then isolated from the matrices of the TCGA patient cohorts shown in Fig. 6. The submatrices thus created were analyzed using the singular value decomposition to generate scores for each sample. Rank listing of samples according to score and sectioning into quartiles based upon those scores and comparison with JAMMIT solution-derived clustering produced the results shown.

### Routine statistical calculations

Results of Fisher’s exact test were produced using the bioinformatic tool online at https://www.scistat.com/statisticaltests/fisher.php. Kaplan-Meier plots were produced using the online statistical tool at https://astatsa.com/LogRankTest/. Perl (Indigoperl build for Microsoft Windows, available through www.indigostar.com) was used primarily for sorting data and performing other non-mathematical operations not readily adapted to use of Excel.

**Table 1.**
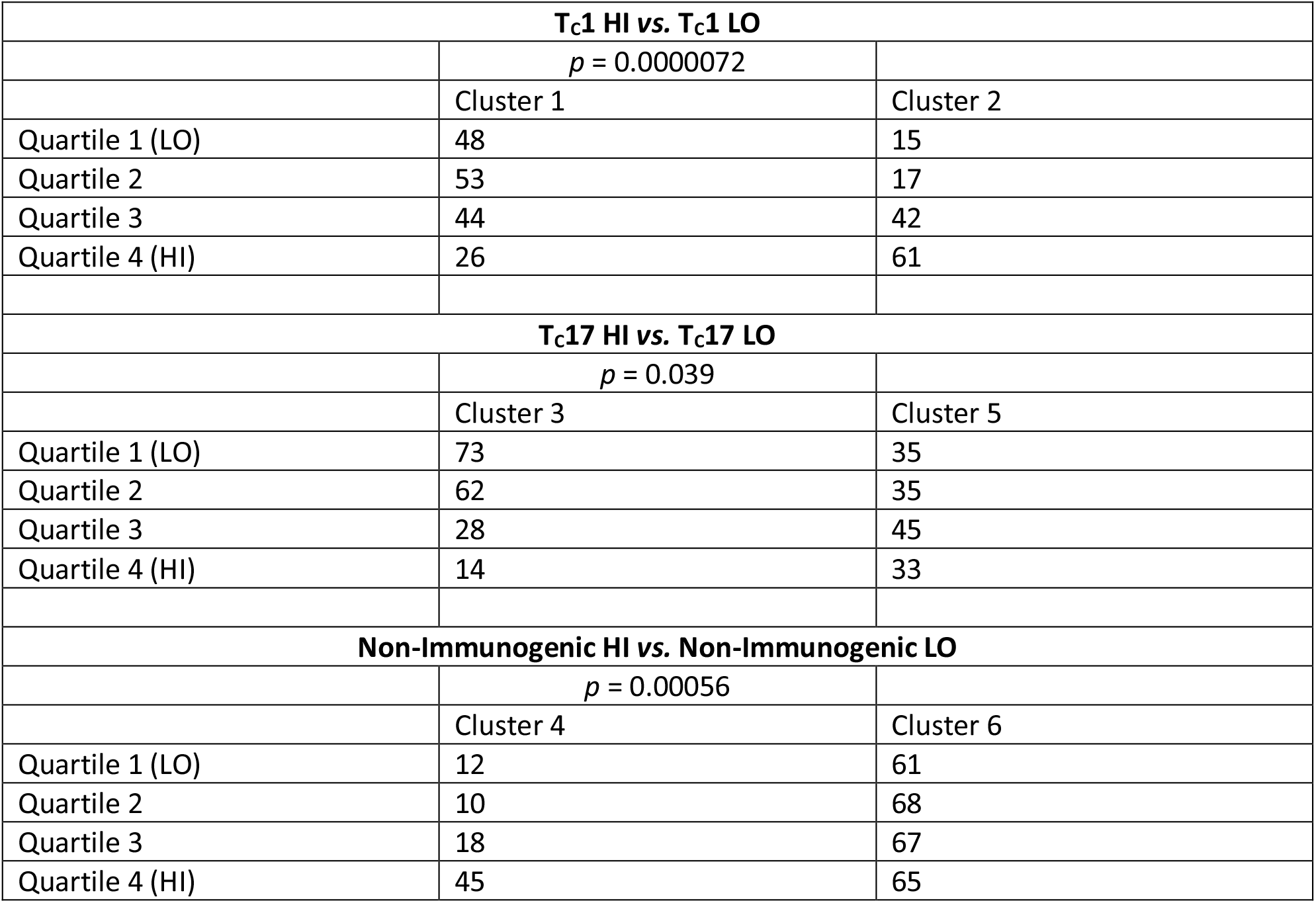
Rank-order expression of the “Warburg” signature in immunophenotypic subclusters for BRCA. The number of samples per quartile in the 6 immunophenotypic subclusters depicted in Figure S3 is at varying levels of the “Warburg effect” as determined by a 3-gene signature. Note that with increasing levels of *PKM*, *SLC16A3* and *SLC2A1*, the relative distribution of samples increases preferentially in one subcluster regardless of immunophenotype, indicating a secondary source of variability in the global transcriptome commensurate with infiltration of tumor by agranulocytes. The *p* value shown indicates the statistical measure of significance among the 2 subclusters in each of the 3 main immunophenotypes at high and low levels of the measured Warburg effect according to Fisher’s exact test.

## Data availability

Data used in all analyses described here are available through the Genomic Data Commons as part of the TCGA project. DbGap data and other data used in validation shown in Fig. 6 are available as referenced below.

## Acknowledgements

This work was supported in part by NIH grants R01CA223490 and R01 CA230514 to Youping Deng and 5P30GM114737, P20GM103466, 5U54MD007601 and 5P30CA071789.

## References

1 Jiao, S., Subudhi, S.K., Aparicio, A., Ge, Z., Guan, B., Miura, Y., Sharma, P., 2019. Differences in Tumor Microenvironment Dictate T Helper Lineage Polarization and Response to Immune Checkpoint Therapy. Cell 179, 1177–1190.e13. https://doi.org/10.1016/j.cell.2019.10.029

2 Blank, C.U., Rozeman, E.A., Fanchi, L.F., Sikorska, K., van de Wiel, B., Kvistborg, P., Krijgsman, O., van den Braber, M., Philips, D., Broeks, A., van Thienen, J.V., Mallo, H.A., Adriaansz, S., ter Meulen, S., Pronk, L.M., Grijpink-Ongering, L.G., Bruining, A., Gittelman, R.M., Warren, S., van Tinteren, H., Peeper, D.S., Haanen, J.B.A.G., van Akkooi, A.C.J., Schumacher, T.N., 2018. Neoadjuvant versus adjuvant ipilimumab plus nivolumab in macroscopic stage III melanoma. Nat Med 24, 1655–1661. https://doi.org/10.1038/s41591-018-0198-0

3 Tsai, J.-P., Lee, M.-H., Hsu, S.-C., Chen, M.-Y., Liu, S.-J., Chang, J.T., Liao, C.-T., Cheng, A.-J., Chong, P., Chu, C.-L., Shen, C.-R., Chen, H.-W., 2012. CD4 + T Cells Disarm or Delete Cytotoxic T Lymphocytes under IL-17–Polarizing Conditions. J.I. 189, 1671–1679. https://doi.org/10.4049/jimmunol.1103447

4 Sheng, I.Y., Diaz-Montero, C.M., Rayman, P., Wei, W., Finke, J.H., Kim, J.S., Pavicic, P.G., Lamenza, M., Company, D., Stephenson, A., Campbell, S., Haber, G., Lee, B., Mian, O., Gilligan, T.D., Rini, B.I., Garcia, J.A., Grivas, P., Ornstein, M.C., 2020. Blood Myeloid-Derived Suppressor Cells Correlate with Neutrophil-to-Lymphocyte Ratio and Overall Survival in Metastatic Urothelial Carcinoma. Targeted Oncology 15, 211–220. https://doi.org/10.1007/s11523-020-00707-z

5 Zhang, M., He, Y., Sun, X., Li, Q., Wang, W., Zhao, A., Di, W., 2014. A high M1/M2 ratio of tumor-associated macrophages is associated with extended survival in ovarian cancer patients. J Ovarian Res 7, 19. https://doi.org/10.1186/1757-2215-7-19

6 Deng, Y., Cheng, J., Fu, B., Liu, W., Chen, G., Zhang, Q., Yang, Y., 2017. Hepatic carcinoma-associated fibroblasts enhance immune suppression by facilitating the generation of myeloid-derived suppressor cells. Oncogene 36, 1090–1101. https://doi.org/10.1038/onc.2016.273

7 Edin, S., Wikberg, M.L., Dahlin, A.M., Rutegård, J., Öberg, Å., Oldenborg, P.-A., Palmqvist, R., 2012. The Distribution of Macrophages with a M1 or M2 Phenotype in Relation to Prognosis and the Molecular Characteristics of Colorectal Cancer. PLoS ONE 7, e47045. https://doi.org/10.1371/journal.pone.0047045

8 Hu, H., Hang, J.-J., Han, T., Zhuo, M., Jiao, F., Wang, L.-W., 2016. The M2 phenotype of tumor-associated macrophages in the stroma confers a poor prognosis in pancreatic cancer. Tumor Biol. 37, 8657–8664. https://doi.org/10.1007/s13277-015-4741-z

9 The Cancer Genome Atlas Research Network, Weinstein, J.N., Collisson, E.A., Mills, G.B., Shaw, K.R.M., Ozenberger, B.A., Ellrott, K., Shmulevich, I., Sander, C., Stuart, J.M., 2013. The Cancer Genome Atlas Pan-Cancer analysis project. Nat Genet 45, 1113–1120. https://doi.org/10.1038/ng.2764

10 Akbani, R., Akdemir, K.C., Aksoy, B.A., Albert, M., Ally, A., Amin, S.B., Arachchi, H., Arora, A., Auman, J.T., Ayala, B., Baboud, J., Balasundaram, M., Balu, S., Barnabas, N., Bartlett, J., Bartlett, P., Bastian, B.C., Baylin, S.B., Behera, M., Belyaev, D., Benz, C., Bernard, B., Beroukhim, R., Bir, N., Black, A.D., Bodenheimer, T., Boice, L., Boland, G.M., Bono, R., Bootwalla, M.S., Bosenberg, M., Bowen, J., Bowlby, R., Bristow, C.A., Brockway-Lunardi, L., Brooks, D., Brzezinski, J., Bshara, W., Buda, E., Burns, W.R., Butterfield, Y.S.N., Button, M., Calderone, T., Cappellini, G.A., Carter, C., Carter, S.L., Cherney, L., Cherniack, A.D., Chevalier, A., Chin, L., Cho, J., Cho, R.J., Choi, Y.-L., Chu, A., Chudamani, S., Cibulskis, K., Ciriello, G., Clarke, A., Coons, S., Cope, L., Crain, D., Curley, E., Danilova, L., D’Atri, S., Davidsen, T., Davies, M.A., Delman, K.A., Demchok, J.A., Deng, Q.A., Deribe, Y.L., Dhalla, N., Dhir, R., DiCara, D., Dinikin, M., Dubina, M., Ebrom, J.S., Egea, S., Eley, G., Engel, J., Eschbacher, J.M., Fedosenko, K.V., Felau, I., Fennell, T., Ferguson, M.L., Fisher, S., Flaherty, K.T., Frazer, S., Frick, J., Fulidou, V., Gabriel, S.B., Gao, J., Gardner, J., Garraway, L.A., Gastier-Foster, J.M., Gaudioso, C., Gehlenborg, N., Genovese, G., Gerken, M., Gershenwald, J.E., Getz, G., Gomez-Fernandez, C., Gribbin, T., Grimsby, J., Gross, B., Guin, R., Gutschner, T., Hadjipanayis, A., Halaban, R., Hanf, B., Haussler, D., Haydu, L.E., Hayes, D.N., Hayward, N.K., Heiman, D.I., Herbert, L., Herman, J.G., Hersey, P., Hoadley, K.A., Hodis, E., Holt, R.A., Hoon, D.SB., Hoppough, S., Hoyle, A.P., Huang, F.W., Huang, M., Huang, S., Hutter, C.M., Ibbs, M., Iype, L., Jacobsen, A., Jakrot, V., Janning, A., Jeck, W.R., Jefferys, S.R., Jensen, M.A., Jones, C.D., Jones, S.J.M., Ju, Z., Kakavand, H., Kang, H., Kefford, R.F., Khuri, F.R., Kim, J., Kirkwood, J.M., Klode, J., Korkut, A., Korski, K., Krauthammer, M., Kucherlapati, R., Kwong, L.N., Kycler, W., Ladanyi, M., Lai, P.H., Laird, P.W., Lander, E., Lawrence, M.S., Lazar, A.J., Łaźniak, R., Lee, D., Lee, J.E., Lee, J., Lee, K., Lee, S., Lee, W., Leporowska, E., Leraas, K.M., Li, H.I., Lichtenberg, T.M., Lichtenstein, L., Lin, P., Ling, S., Liu, J., Liu, O., Liu, W., Long, G.V., Lu, Y., Ma, S., Ma, Y., Mackiewicz, A., Mahadeshwar, H.S., Malke, J., Mallery, D., Manikhas, G.M., Mann, G.J., Marra, M.A., Matejka, B., Mayo, M., Mehrabi, S., Meng, S., Meyerson, M., Mieczkowski, P.A., Miller, J.P., Miller, M.L., Mills, G.B., Moiseenko, F., Moore, R.A., Morris, S., Morrison, C., Morton, D., Moschos, S., Mose, L.E., Muller, F.L., Mungall, A.J., Murawa, D., Murawa, P., Murray, B.A., Nezi, L., Ng, S., Nicholson, D., Noble, M.S., Osunkoya, A., Owonikoko, T.K., Ozenberger, B.A., Pagani, E., Paklina, O.V., Pantazi, A., Parfenov, M., Parfitt, J., Park, P.J., Park, W.-Y., Parker, J.S., Passarelli, F., Penny, R., Perou, C.M., Pihl, T.D., Potapova, O., Prieto, V.G., Protopopov, A., Quinn, M.J., Radenbaugh, A., Rai, K., Ramalingam, S.S., Raman, A.T., Ramirez, N.C., Ramirez, R., Rao, U., Rathmell, W.K., Ren, X., Reynolds, S.M., Roach, J., Robertson, A.G., Ross, M.I., Roszik, J., Russo, G., Saksena, G., Saller, C., Samuels, Y., Sander, Chris, Sander, Cindy, Sandusky, G., Santoso, N., Saul, M., Saw, R.PM., Schadendorf, D., Schein, J.E., Schultz, N., Schumacher, S.E., Schwallier, C., Scolyer, R.A., Seidman, J., Sekhar, P.C., Sekhon, H.S., Senbabaoglu, Y., Seth, S., Shannon, K.F., Sharpe, S., Sharpless, N.E., Shaw, K.R.M., Shelton, C., Shelton, T., Shen, R., Sheth, M., Shi, Y., Shiau, C.J., Shmulevich, I., Sica, G.L., Simons, J.V., Sinha, R., Sipahimalani, P., Sofia, H.J., Soloway, M.G., Song, X., Sougnez, C., Spillane, A.J., Spychała, A., Stretch, J.R., Stuart, J., Suchorska, W.M., Sucker, A., Sumer, S.O., Sun, Y., Synott, M., Tabak, B., Tabler, T.R., Tam, A., Tan, D., Tang, J., Tarnuzzer, R., Tarvin, K., Tatka, H., Taylor, B.S., Teresiak, M., Thiessen, N., Thompson, J.F., Thorne, L., Thorsson, V., Trent, J.M., Triche, T.J., Tsai, K.Y., Tsou, P., Van Den Berg, D.J., Van Allen, E.M., Veluvolu, U., Verhaak, R.G., Voet, D., Voronina, O., Walter, V., Walton, J.S., Wan, Y., Wang, Y., Wang, Z., Waring, S., Watson, I.R., Weinhold, N., Weinstein, J.N., Weisenberger, D.J., White, P., Wilkerson, M.D., Wilmott, J.S., Wise, L., Wiznerowicz, M., Woodman, S.E., Wu, C.-J., Wu, C.-C., Wu, J., Wu, Y., Xi, R., Xu, A.W., Yang, D., Yang, Liming, Yang, Lixing, Zack, T.I., Zenklusen, J.C., Zhang, H., Zhang, J., Zhang, W., Zhao, X., Zhu, J., Zhu, K., Zimmer, L., Zmuda, E., Zou, L., 2015. Genomic Classification of Cutaneous Melanoma. Cell 161, 1681–1696. https://doi.org/10.1016/j.cell.2015.05.044

11 Ally, A., Balasundaram, M., Carlsen, R., Chuah, E., Clarke, A., Dhalla, N., Holt, R.A., Jones, S.J.M., Lee, D., Ma, Y., Marra, M.A., Mayo, M., Moore, R.A., Mungall, A.J., Schein, J.E., Sipahimalani, P., Tam, A., Thiessen, N., Cheung, D., Wong, T., Brooks, D., Robertson, A.G., Bowlby, R., Mungall, K., Sadeghi, S., Xi, L., Covington, K., Shinbrot, E., Wheeler, D.A., Gibbs, R.A., Donehower, L.A., Wang, L., Bowen, J., Gastier-Foster, J.M., Gerken, M., Helsel, C., Leraas, K.M., Lichtenberg, T.M., Ramirez, N.C., Wise, L., Zmuda, E., Gabriel, S.B., Meyerson, M., Cibulskis, C., Murray, B.A., Shih, J., Beroukhim, R., Cherniack, A.D., Schumacher, S.E., Saksena, G., Pedamallu, C.S., Chin, L., Getz, G., Noble, M., Zhang, Hailei, Heiman, D., Cho, J., Gehlenborg, N., Saksena, G., Voet, D., Lin, P., Frazer, S., Defreitas, T., Meier, S., Lawrence, M., Kim, J., Creighton, C.J., Muzny, D., Doddapaneni, H., Hu, J., Wang, M., Morton, D., Korchina, V., Han, Y., Dinh, H., Lewis, L., Bellair, M., Liu, X., Santibanez, J., Glenn, R., Lee, S., Hale, W., Parker, J.S., Wilkerson, M.D., Hayes, D.N., Reynolds, S.M., Shmulevich, I., Zhang, W., Liu, Y., Iype, L., Makhlouf, H., Torbenson, M.S., Kakar, S., Yeh, M.M., Jain, D., Kleiner, D.E., Jain, D., Dhanasekaran, R., El-Serag, H.B., Yim, S.Y., Weinstein, J.N., Mishra, L., Zhang, Jianping, Akbani, R., Ling, S., Ju, Z., Su, X., Hegde, A.M., Mills, G.B., Lu, Y., Chen, J., Lee, J.-S., Sohn, B.H., Shim, J.J., Tong, P., Aburatani, H., Yamamoto, S., Tatsuno, K., Li, W., Xia, Z., Stransky, N., Seiser, E., Innocenti, F., Gao, J., Kundra, R., Zhang, Hongxin, Heins, Z., Ochoa, A., Sander, C., Ladanyi, M., Shen, R., Arora, A., Sanchez-Vega, F., Schultz, N., Kasaian, K., Radenbaugh, A., Bissig, K.- D., Moore, D.D., Totoki, Y., Nakamura, H., Shibata, T., Yau, C., Graim, K., Stuart, J., Haussler, D., Slagle, B.L., Ojesina, A.I., Katsonis, P., Koire, A., Lichtarge, O., Hsu, T.-K., Ferguson, M.L., Demchok, J.A., Felau, I., Sheth, M., Tarnuzzer, R., Wang, Z., Yang, L., Zenklusen, J.C., Zhang, Jiashan, Hutter, C.M., Sofia, H.J., Verhaak, R.G.W., Zheng, S., Lang, F., Chudamani, S., Liu, J., Lolla, L., Wu, Y., Naresh, R., Pihl, T., Sun, C., Wan, Y., Benz, C., Perou, A.H., Thorne, L.B., Boice, L., Huang, M., Rathmell, W.K., Noushmehr, H., Saggioro, F.P., Tirapelli, D.P. da C., Junior, C.G.C., Mente, E.D., Silva, O. de C., Trevisan, F.A., Kang, K.J., Ahn, K.S., Giama, N.H., Moser, C.D., Giordano, T.J., Vinco, M., Welling, T.H., Crain, D., Curley, E., Gardner, J., Mallery, D., Morris, S., Paulauskis, J., Penny, R., Shelton, C., Shelton, T., Kelley, R., Park, J.-W., Chandan, V.S., Roberts, L.R., Bathe, O.F., Hagedorn, C.H., Auman, J.T., O’Brien, D.R., Kocher, J.-P.A., Jones, C.D., Mieczkowski, P.A., Perou, C.M., Skelly, T., Tan, D., Veluvolu, U., Balu, S., Bodenheimer, T., Hoyle, A.P., Jefferys, S.R., Meng, S., Mose, L.E., Shi, Y., Simons, J.V., Soloway, M.G., Roach, J., Hoadley, K.A., Baylin, S.B., Shen, H., Hinoue, T., Bootwalla, M.S., Van Den Berg, D.J., Weisenberger, D.J., Lai, P.H., Holbrook, A., Berrios, M., Laird, P.W., 2017. Comprehensive and Integrative Genomic Characterization of Hepatocellular Carcinoma. Cell 169, 1327–1341.e23. https://doi.org/10.1016/j.cell.2017.05.046

12 The ICGC/TCGA Pan-Cancer Analysis of Whole Genomes Consortium, 2020. Pan-cancer analysis of whole genomes. Nature 578, 82–93. https://doi.org/10.1038/s41586-020-1969-6

13 Hoadley, K.A., Yau, C., Hinoue, T., Wolf, D.M., Lazar, A.J., Drill, E., Shen, R., Taylor, A.M., Cherniack, A.D., Thorsson, Vésteinn, Akbani, R., Bowlby, R., Wong, C.K., Wiznerowicz, M., Sanchez-Vega, F., Robertson, A.G., Schneider, B.G., Lawrence, M.S., Noushmehr, H., Malta, T.M., Stuart, J.M., Benz, C.C., Laird, P.W., Caesar-Johnson, S.J., Demchok, J.A., Felau, I., Kasapi, M., Ferguson, M.L., Hutter, C.M., Sofia, H.J., Tarnuzzer, R., Wang, Z., Yang, L., Zenklusen, J.C., Zhang, J. (Julia), Chudamani, S., Liu, J., Lolla, L., Naresh, R., Pihl, T., Sun, Q., Wan, Y., Wu, Y., Cho, J., DeFreitas, T., Frazer, S., Gehlenborg, N., Getz, G., Heiman, D.I., Kim, J., Lawrence, M.S., Lin, P., Meier, S., Noble, M.S., Saksena, G., Voet, D., Zhang, Hailei, Bernard, B., Chambwe, N., Dhankani, V., Knijnenburg, T., Kramer, R., Leinonen, K., Liu, Y., Miller, M., Reynolds, S., Shmulevich, I., Thorsson, Vesteinn, Zhang, W., Akbani, R., Broom, B.M., Hegde, A.M., Ju, Z., Kanchi, R.S., Korkut, A., Li, J., Liang, H., Ling, S., Liu, W., Lu, Y., Mills, G.B., Ng, K.-S., Rao, A., Ryan, M., Wang, Jing, Weinstein, J.N., Zhang, J., Abeshouse, A., Armenia, J., Chakravarty, D., Chatila, W.K., de Bruijn, I., Gao, J., Gross, B.E., Heins, Z.J., Kundra, R., La, K., Ladanyi, M., Luna, A., Nissan, M.G., Ochoa, A., Phillips, S.M., Reznik, E., Sanchez-Vega, F., Sander, C., Schultz, N., Sheridan, R., Sumer, S.O., Sun, Y., Taylor, B.S., Wang, Jioajiao, Zhang, Hongxin, Anur, P., Peto, M., Spellman, P., Benz, C., Stuart, J.M., Wong, C.K., Yau, C., Hayes, D.N., Parker, J.S., Wilkerson, M.D., Ally, A., Balasundaram, M., Bowlby, R., Brooks, D., Carlsen, R., Chuah, E., Dhalla, N., Holt, R., Jones, S.J.M., Kasaian, K., Lee, D., Ma, Y., Marra, M.A., Mayo, M., Moore, R.A., Mungall, A.J., Mungall, K., Robertson, A.G., Sadeghi, S., Schein, J.E., Sipahimalani, P., Tam, A., Thiessen, N., Tse, K., Wong, T., Berger, A.C., Beroukhim, R., Cherniack, A.D., Cibulskis, C., Gabriel, S.B., Gao, G.F., Ha, G., Meyerson, M., Schumacher, S.E., Shih, J., Kucherlapati, M.H., Kucherlapati, R.S., Baylin, S., Cope, L., Danilova, L., Bootwalla, M.S., Lai, P.H., Maglinte, D.T., Van Den Berg, D.J., Weisenberger, D.J., Auman, J.T., Balu, S., Bodenheimer, T., Fan, C., Hoadley, K.A., Hoyle, A.P., Jefferys, S.R., Jones, C.D., Meng, S., Mieczkowski, P.A., Mose, L.E., Perou, A.H., Perou, C.M., Roach, J., Shi, Y., Simons, J.V., Skelly, T., Soloway, M.G., Tan, D., Veluvolu, U., Fan, H., Hinoue, T., Laird, P.W., Shen, H., Zhou, W., Bellair, M., Chang, K., Covington, K., Creighton, C.J., Dinh, H., Doddapaneni, H., Donehower, L.A., Drummond, J., Gibbs, R.A., Glenn, R., Hale, W., Han, Y., Hu, J., Korchina, V., Lee, S., Lewis, L., Li, W., Liu, X., Morgan, M., Morton, D., Muzny, D., Santibanez, J., Sheth, M., Shinbrot, E., Wang, L., Wang, M., Wheeler, D.A., Xi, L., Zhao, F., Hess, J., Appelbaum, E.L., Bailey, M., Cordes, M.G., Ding, L., Fronick, C.C., Fulton, L.A., Fulton, R.S., Kandoth, C., Mardis, E.R., McLellan, M.D., Miller, C.A., Schmidt, H.K., Wilson, R.K., Crain, D., Curley, E., Gardner, J., Lau, K., Mallery, D., Morris, S., Paulauskis, J., Penny, R., Shelton, C., Shelton, T., Sherman, M., Thompson, E., Yena, P., Bowen, J., Gastier-Foster, J.M., Gerken, M., Leraas, K.M., Lichtenberg, T.M., Ramirez, N.C., Wise, L., Zmuda, E., Corcoran, N., Costello, T., Hovens, C., Carvalho, A.L., de Carvalho, A.C., Fregnani, J.H., Longatto-Filho, A., Reis, R.M., Scapulatempo-Neto, C., Silveira, H.C.S., Vidal, D.O., Burnette, A., Eschbacher, J., Hermes, B., Noss, A., Singh, R., Anderson, M.L., Castro, P.D., Ittmann, M., Huntsman, D., Kohl, B., Le, X., Thorp, R., Andry, C., Duffy, E.R., Lyadov, V., Paklina, O., Setdikova, G., Shabunin, A., Tavobilov, M., McPherson, C., Warnick, R., Berkowitz, R., Cramer, D., Feltmate, C., Horowitz, N., Kibel, A., Muto, M., Raut, C.P., Malykh, A., Barnholtz-Sloan, J.S., Barrett, W., Devine, K., Fulop, J., Ostrom, Q.T., Shimmel, K., Wolinsky, Y., Sloan, A.E., De Rose, A., Giuliante, F., Goodman, M., Karlan, B.Y., Hagedorn, C.H., Eckman, J., Harr, J., Myers, J., Tucker, K., Zach, L.A., Deyarmin, B., Hu, H., Kvecher, L., Larson, C., Mural, R.J., Somiari, S., Vicha, A., Zelinka, T., Bennett, J., Iacocca, M., Rabeno, B., Swanson, P., Latour, M., Lacombe, L., Têtu, B., Bergeron, A., McGraw, M., Staugaitis, S.M., Chabot, J., Hibshoosh, H., Sepulveda, A., Su, T., Wang, T., Potapova, O., Voronina, O., Desjardins, L., Mariani, O., Roman-Roman, S., Sastre, X., Stern, M.-H., Cheng, F., Signoretti, S., Berchuck, A., Bigner, D., Lipp, E., Marks, J., McCall, S., McLendon, R., Secord, A., Sharp, A., Behera, M., Brat, D.J., Chen, A., Delman, K., Force, S., Khuri, F., Magliocca, K., Maithel, S., Olson, J.J., Owonikoko, T., Pickens, A., Ramalingam, S., Shin, D.M., Sica, G., Van Meir, E.G., Zhang, Hongzheng, Eijckenboom, W., Gillis, A., Korpershoek, E., Looijenga, L., Oosterhuis, W., Stoop, H., van Kessel, K.E., Zwarthoff, E.C., Calatozzolo, C., Cuppini, L., Cuzzubbo, S., DiMeco, F., Finocchiaro, G., Mattei, L., Perin, A., Pollo, B., Chen, C., Houck, J., Lohavanichbutr, P., Hartmann, A., Stoehr, C., Stoehr, R., Taubert, H., Wach, S., Wullich, B., Kycler, W., Murawa, D., Wiznerowicz, M., Chung, K., Edenfield, W.J., Martin, J., Baudin, E., Bubley, G., Bueno, R., De Rienzo, A., Richards, W.G., Kalkanis, S., Mikkelsen, T., Noushmehr, H., Scarpace, L., Girard, N., Aymerich, M., Campo, E., Giné, E., Guillermo, A.L., Van Bang, N., Hanh, P.T., Phu, B.D., Tang, Y., Colman, H., Evason, K., Dottino, P.R., Martignetti, J.A., Gabra, H., Juhl, H., Akeredolu, T., Stepa, S., Hoon, D., Ahn, K., Kang, K.J., Beuschlein, F., Breggia, A., Birrer, M., Bell, D., Borad, M., Bryce, A.H., Castle, E., Chandan, V., Cheville, J., Copland, J.A., Farnell, M., Flotte, T., Giama, N., Ho, T., Kendrick, M., Kocher, J.-P., Kopp, K., Moser, C., Nagorney, D., O’Brien, D., O’Neill, B.P., Patel, T., Petersen, G., Que, F., Rivera, M., Roberts, L., Smallridge, R., Smyrk, T., Stanton, M., Thompson, R.H., Torbenson, M., Yang, J.D., Zhang, L., Brimo, F., Ajani, J.A., Gonzalez, A.M.A., Behrens, C., Bondaruk, olanta, Broaddus, R., Czerniak, B., Esmaeli, B., Fujimoto, J., Gershenwald, J., Guo, C., Lazar, A.J., Logothetis, C., Meric-Bernstam, F., Moran, C., Ramondetta, L., Rice, D., Sood, A., Tamboli, P., Thompson, T., Troncoso, P., Tsao, A., Wistuba, I., Carter, C., Haydu, L., Hersey, P., Jakrot, V., Kakavand, H., Kefford, R., Lee, K., Long, G., Mann, G., Quinn, M., Saw, R., Scolyer, R., Shannon, K., Spillane, A., Stretch, J., Synott, M., Thompson, J., Wilmott, J., Al-Ahmadie, H., Chan, T.A., Ghossein, R., Gopalan, A., Levine, D.A., Reuter, V., Singer, S., Singh, B., Tien, N.V., Broudy, T., Mirsaidi, C., Nair, P., Drwiega, P., Miller, J., Smith, J., Zaren, H., Park, J.-W., Hung, N.P., Kebebew, E., Linehan, W.M., Metwalli, A.R., Pacak, K., Pinto, P.A., Schiffman, M., Schmidt, L.S., Vocke, C.D., Wentzensen, N., Worrell, R., Yang, H., Moncrieff, M., Goparaju, C., Melamed, J., Pass, H., Botnariuc, N., Caraman, I., Cernat, M., Chemencedji, I., Clipca, A., Doruc, S., Gorincioi, G., Mura, S., Pirtac, M., Stancul, I., Tcaciuc, D., Albert, M., Alexopoulou, I., Arnaout, A., Bartlett, J., Engel, J., Gilbert, S., Parfitt, J., Sekhon, H., Thomas, G., Rassl, D.M., Rintoul, R.C., Bifulco, C., Tamakawa, R., Urba, W., Hayward, N., Timmers, H., Antenucci, A., Facciolo, F., Grazi, G., Marino, M., Merola, R., de Krijger, R., Gimenez-Roqueplo, A.-P., Piché, A., Chevalier, S., McKercher, G., Birsoy, K., Barnett, G., Brewer, C., Farver, C., Naska, T., Pennell, N.A., Raymond, D., Schilero, C., Smolenski, K., Williams, F., Morrison, C., Borgia, J.A., Liptay, M.J., Pool, M., Seder, C.W., Junker, K., Omberg, L., Dinkin, M., Manikhas, G., Alvaro, D., Bragazzi, M.C., Cardinale, V., Carpino, G., Gaudio, E., Chesla, D., Cottingham, S., Dubina, M., Moiseenko, F., Dhanasekaran, R., Becker, K.-F., Janssen, K.-P., Slotta-Huspenina, J., Abdel-Rahman, M.H., Aziz, D., Bell, S., Cebulla, C.M., Davis, A., Duell, R., Elder, J.B., Hilty, J., Kumar, B., Lang, J., Lehman, N.L., Mandt, R., Nguyen, P., Pilarski, R., Rai, K., Schoenfield, L., Senecal, K., Wakely, P., Hansen, P., Lechan, R., Powers, J., Tischler, A., Grizzle, W.E., Sexton, K.C., Kastl, A., Henderson, J., Porten, S., Waldmann, J., Fassnacht, M., Asa, S.L., Schadendorf, D., Couce, M., Graefen, M., Huland, H., Sauter, G., Schlomm, T., Simon, R., Tennstedt, P., Olabode, O., Nelson, M., Bathe, O., Carroll, P.R., Chan, J.M., Disaia, P., Glenn, P., Kelley, R.K., Landen, C.N., Phillips, J., Prados, M., Simko, J., Smith-McCune, K., VandenBerg, S., Roggin, K., Fehrenbach, A., Kendler, A., Sifri, S., Steele, R., Jimeno, A., Carey, F., Forgie, I., Mannelli, M., Carney, M., Hernandez, B., Campos, B., Herold-Mende, C., Jungk, C., Unterberg, A., von Deimling, A., Bossler, A., Galbraith, J., Jacobus, L., Knudson, M., Knutson, T., Ma, D., Milhem, M., Sigmund, R., Godwin, A.K., Madan, R., Rosenthal, H.G., Adebamowo, C., Adebamowo, S.N., Boussioutas, A., Beer, D., Giordano, T., Mes-Masson, A.-M., Saad, F., Bocklage, T., Landrum, L., Mannel, R., Moore, K., Moxley, K., Postier, R., Walker, J., Zuna, R., Feldman, M., Valdivieso, F., Dhir, R., Luketich, J., Pinero, E.M.M., Quintero-Aguilo, M., Carlotti, C.G., Dos Santos, J.S., Kemp, R., Sankarankuty, A., Tirapelli, D., Catto, J., Agnew, K., Swisher, E., Creaney, J., Robinson, B., Shelley, C.S., Godwin, E.M., Kendall, S., Shipman, C., Bradford, C., Carey, T., Haddad, A., Moyer, J., Peterson, L., Prince, M., Rozek, L., Wolf, G., Bowman, R., Fong, K.M., Yang, I., Korst, R., Rathmell, W.K., Fantacone-Campbell, J.L., Hooke, J.A., Kovatich, A.J., Shriver, C.D., DiPersio, J., Drake, B., Govindan, R., Heath, S., Ley, T., Van Tine, B., Westervelt, P., Rubin, M.A., Lee, J.I., Aredes, N.D., Mariamidze, A., 2018. Cell-of-Origin Patterns Dominate the Molecular Classification of 10,000 Tumors from 33 Types of Cancer. Cell 173, 291–304.e6. https://doi.org/10.1016/j.cell.2018.03.022

14 Lizotte, P.H., Ivanova, E.V., Awad, M.M., Jones, R.E., Keogh, L., Liu, H., Dries, R., Almonte, C., Herter-Sprie, G.S., Santos, A., Feeney, N.B., Paweletz, C.P., Kulkarni, M.M., Bass, A.J., Rustgi, A.K., Yuan, G.-C., Kufe, D.W., Jänne, P.A., Hammerman, P.S., Sholl, L.M., Hodi, F.S., Richards, W.G., Bueno, R., English, J.M., Bittinger, M.A., Wong, K.-K., 2016. Multiparametric profiling of non–small-cell lung cancers reveals distinct immunophenotypes. JCI Insight 1. https://doi.org/10.1172/jci.insight.89014

15 Zhou, Y.-J., Zhu, G.-Q., Lu, X.-F., Zheng, K.I., Wang, Q.-W., Chen, J.-N., Zhang, Q.-W., Yan, F.-R., Li, X.-B., 2020. Identification and validation of tumour microenvironment-based immune molecular subgroups for gastric cancer: immunotherapeutic implications. Cancer Immunol Immunother 69, 1057–1069. https://doi.org/10.1007/s00262-020-02525-8

16 Okimoto, G., Zeinalzadeh, A., Wenska, T., Loomis, M., Nation, J.B., Fabre, T., Tiirikainen, M., Hernandez, B., Chan, O., Wong, L., Kwee, S., 2016. Joint analysis of multiple high-dimensional data types using sparse matrix approximations of rank-1 with applications to ovarian and liver cancer. BioData Min 9, 24–24. https://doi.org/10.1186/s13040-016-0103-7

17 Kwee, S.A., Okimoto, G.S., Chan, O.T., Tiirikainen, M., Wong, L.L., n.d. Metabolic characteristics distinguishing intrahepatic cholangiocarcinoma: a negative pilot study of 18F-fluorocholine PET/CT clarified by transcriptomic analysis 11.

18 Nagarsheth, N., Wicha, M.S., Zou, W., 2017. Chemokines in the cancer microenvironment and their relevance in cancer immunotherapy. Nat Rev Immunol 17, 559–572. https://doi.org/10.1038/nri.2017.49

19 Craig, Chandler, Gellert, Lambowitz, Rice, Sandmeyer (Eds.), 2015. V(D)J Recombination: Mechanism, Errors, and Fidelity, in: Mobile DNA III. American Society of Microbiology, pp. 313–324. https://doi.org/10.1128/microbiolspec.MDNA3-0041-2014

20 Furness, A.J.S., Vargas, F.A., Peggs, K.S., Quezada, S.A., 2014. Impact of tumour microenvironment and Fc receptors on the activity of immunomodulatory antibodies. Trends in Immunology 35, 290–298. https://doi.org/10.1016/j.it.2014.05.002

21 Alexander Lex, Nils Gehlenborg, Hendrik Strobelt, Romain Vuillemot, Hanspeter Pfister, UpSet: Visualization of Intersecting Sets, IEEE Transactions on Visualization and Computer Graphics (InfoVis ’14), vol. 20, no. 12, pp. 1983–1992, 2014. doi:10.1109/TVCG.2014.2346248

22 Li, S., Morita, H., Sokolowska, M., Tan, G., Boonpiyathad, T., Opitz, L., Orimo, K., Archer, S.K., Jansen, K., Tang, M.L.K., Purcell, D., Plebanski, M., Akdis, C.A., 2019. Gene expression signatures of circulating human type 1, 2, and 3 innate lymphoid cells. Journal of Allergy and Clinical Immunology 143, 2321–2325. https://doi.org/10.1016/j.jaci.2019.01.047

23 Annunziato, F., Romagnani, C., Romagnani, S., 2015. The 3 major types of innate and adaptive cell-mediated effector immunity. Journal of Allergy and Clinical Immunology 135, 626–635. https://doi.org/10.1016/j.jaci.2014.11.001

24 Thorsson, Vésteinn, Gibbs, D.L., Brown, S.D., Wolf, D., Bortone, D.S., Ou Yang, T.-H., Porta-Pardo, E., Gao, G.F., Plaisier, C.L., Eddy, J.A., Ziv, E., Culhane, A.C., Paull, E.O., Sivakumar, I.K.A., Gentles, A.J., Malhotra, R., Farshidfar, F., Colaprico, A., Parker, J.S., Mose, L.E., Vo, N.S., Liu, Jianfang, Liu, Y., Rader, J., Dhankani, V., Reynolds, S.M., Bowlby, R., Califano, A., Cherniack, A.D., Anastassiou, D., Bedognetti, D., Mokrab, Y., Newman, A.M., Rao, A., Chen, K., Krasnitz, A., Hu, H., Malta, T.M., Noushmehr, H., Pedamallu, C.S., Bullman, S., Ojesina, A.I., Lamb, A., Zhou, W., Shen, H., Choueiri, T.K., Weinstein, J.N., Guinney, J., Saltz, J., Holt, R.A., Rabkin, C.S., Lazar, A.J., Serody, J.S., Demicco, E.G., Disis, M.L., Vincent, B.G., Shmulevich, I., Caesar-Johnson, S.J., Demchok, J.A., Felau, I., Kasapi, M., Ferguson, M.L., Hutter, C.M., Sofia, H.J., Tarnuzzer, R., Wang, Z., Yang, L., Zenklusen, J.C., Zhang, J. (Julia), Chudamani, S., Liu, Jia, Lolla, L., Naresh, R., Pihl, T., Sun, Q., Wan, Y., Wu, Y., Cho, J., DeFreitas, T., Frazer, S., Gehlenborg, N., Getz, G., Heiman, D.I., Kim, J., Lawrence, M.S., Lin, P., Meier, S., Noble, M.S., Saksena, G., Voet, D., Zhang, Hailei, Bernard, B., Chambwe, N., Dhankani, V., Knijnenburg, T., Kramer, R., Leinonen, K., Liu, Y., Miller, M., Reynolds, S., Shmulevich, I., Thorsson, Vesteinn, Zhang, W., Akbani, R., Broom, B.M., Hegde, A.M., Ju, Z., Kanchi, R.S., Korkut, A., Li, J., Liang, H., Ling, S., Liu, W., Lu, Y., Mills, G.B., Ng, K.-S., Rao, A., Ryan, M., Wang, Jing, Weinstein, J.N., Zhang, J., Abeshouse, A., Armenia, J., Chakravarty, D., Chatila, W.K., de Bruijn, I., Gao, J., Gross, B.E., Heins, Z.J., Kundra, R., La, K., Ladanyi, M., Luna, A., Nissan, M.G., Ochoa, A., Phillips, S.M., Reznik, E., Sanchez-Vega, F., Sander, C., Schultz, N., Sheridan, R., Sumer, S.O., Sun, Y., Taylor, B.S., Wang, Jioajiao, Zhang, Hongxin, Anur, P., Peto, M., Spellman, P., Benz, C., Stuart, J.M., Wong, C.K., Yau, C., Hayes, D.N., Parker, J.S., Wilkerson, M.D., Ally, A., Balasundaram, M., Bowlby, R., Brooks, D., Carlsen, R., Chuah, E., Dhalla, N., Holt, R., Jones, S.J.M., Kasaian, K., Lee, D., Ma, Y., Marra, M.A., Mayo, M., Moore, R.A., Mungall, A.J., Mungall, K., Robertson, A.G., Sadeghi, S., Schein, J.E., Sipahimalani, P., Tam, A., Thiessen, N., Tse, K., Wong, T., Berger, A.C., Beroukhim, R., Cherniack, A.D., Cibulskis, C., Gabriel, S.B., Gao, G.F., Ha, G., Meyerson, M., Schumacher, S.E., Shih, J., Kucherlapati, M.H., Kucherlapati, R.S., Baylin, S., Cope, L., Danilova, L., Bootwalla, M.S., Lai, P.H., Maglinte, D.T., Van Den Berg, D.J., Weisenberger, D.J., Auman, J.T., Balu, S., Bodenheimer, T., Fan, C., Hoadley, K.A., Hoyle, A.P., Jefferys, S.R., Jones, C.D., Meng, S., Mieczkowski, P.A., Mose, L.E., Perou, A.H., Perou, C.M., Roach, J., Shi, Y., Simons, J.V., Skelly, T., Soloway, M.G., Tan, D., Veluvolu, U., Fan, H., Hinoue, T., Laird, P.W., Shen, H., Zhou, W., Bellair, M., Chang, K., Covington, K., Creighton, C.J., Dinh, H., Doddapaneni, H., Donehower, L.A., Drummond, J., Gibbs, R.A., Glenn, R., Hale, W., Han, Y., Hu, J., Korchina, V., Lee, S., Lewis, L., Li, W., Liu, X., Morgan, M., Morton, D., Muzny, D., Santibanez, J., Sheth, M., Shinbrot, E., Wang, L., Wang, M., Wheeler, D.A., Xi, L., Zhao, F., Hess, J., Appelbaum, E.L., Bailey, M., Cordes, M.G., Ding, L., Fronick, C.C., Fulton, L.A., Fulton, R.S., Kandoth, C., Mardis, E.R., McLellan, M.D., Miller, C.A., Schmidt, H.K., Wilson, R.K., Crain, D., Curley, E., Gardner, J., Lau, K., Mallery, D., Morris, S., Paulauskis, J., Penny, R., Shelton, C., Shelton, T., Sherman, M., Thompson, E., Yena, P., Bowen, J., Gastier-Foster, J.M., Gerken, M., Leraas, K.M., Lichtenberg, T.M., Ramirez, N.C., Wise, L., Zmuda, E., Corcoran, N., Costello, T., Hovens, C., Carvalho, A.L., de Carvalho, A.C., Fregnani, J.H., Longatto-Filho, A., Reis, R.M., Scapulatempo-Neto, C., Silveira, H.C.S., Vidal, D.O., Burnette, A., Eschbacher, J., Hermes, B., Noss, A., Singh, R., Anderson, M.L., Castro, P.D., Ittmann, M., Huntsman, D., Kohl, B., Le, X., Thorp, R., Andry, C., Duffy, E.R., Lyadov, V., Paklina, O., Setdikova, G., Shabunin, A., Tavobilov, M., McPherson, C., Warnick, R., Berkowitz, R., Cramer, D., Feltmate, C., Horowitz, N., Kibel, A., Muto, M., Raut, C.P., Malykh, A., Barnholtz-Sloan, J.S., Barrett, W., Devine, K., Fulop, J., Ostrom, Q.T., Shimmel, K., Wolinsky, Y., Sloan, A.E., De Rose, A., Giuliante, F., Goodman, M., Karlan, B.Y., Hagedorn, C.H., Eckman, J., Harr, J., Myers, J., Tucker, K., Zach, L.A., Deyarmin, B., Hu, H., Kvecher, L., Larson, C., Mural, R.J., Somiari, S., Vicha, A., Zelinka, T., Bennett, J., Iacocca, M., Rabeno, B., Swanson, P., Latour, M., Lacombe, L., Têtu, B., Bergeron, A., McGraw, M., Staugaitis, S.M., Chabot, J., Hibshoosh, H., Sepulveda, A., Su, T., Wang, T., Potapova, O., Voronina, O., Desjardins, L., Mariani, O., Roman-Roman, S., Sastre, X., Stern, M.-H., Cheng, F., Signoretti, S., Berchuck, A., Bigner, D., Lipp, E., Marks, J., McCall, S., McLendon, R., Secord, A., Sharp, A., Behera, M., Brat, D.J., Chen, A., Delman, K., Force, S., Khuri, F., Magliocca, K., Maithel, S., Olson, J.J., Owonikoko, T., Pickens, A., Ramalingam, S., Shin, D.M., Sica, G., Van Meir, E.G., Zhang, Hongzheng, Eijckenboom, W., Gillis, A., Korpershoek, E., Looijenga, L., Oosterhuis, W., Stoop, H., van Kessel, K.E., Zwarthoff, E.C., Calatozzolo, C., Cuppini, L., Cuzzubbo, S., DiMeco, F., Finocchiaro, G., Mattei, L., Perin, A., Pollo, B., Chen, C., Houck, J., Lohavanichbutr, P., Hartmann, A., Stoehr, C., Stoehr, R., Taubert, H., Wach, S., Wullich, B., Kycler, W., Murawa, D., Wiznerowicz, M., Chung, K., Edenfield, W.J., Martin, J., Baudin, E., Bubley, G., Bueno, R., De Rienzo, A., Richards, W.G., Kalkanis, S., Mikkelsen, T., Noushmehr, H., Scarpace, L., Girard, N., Aymerich, M., Campo, E., Giné, E., Guillermo, A.L., Van Bang, N., Hanh, P.T., Phu, B.D., Tang, Y., Colman, H., Evason, K., Dottino, P.R., Martignetti, J.A., Gabra, H., Juhl, H., Akeredolu, T., Stepa, S., Hoon, D., Ahn, K., Kang, K.J., Beuschlein, F., Breggia, A., Birrer, M., Bell, D., Borad, M., Bryce, A.H., Castle, E., Chandan, V., Cheville, J., Copland, J.A., Farnell, M., Flotte, T., Giama, N., Ho, T., Kendrick, M., Kocher, J.-P., Kopp, K., Moser, C., Nagorney, D., O’Brien, D., O’Neill, B.P., Patel, T., Petersen, G., Que, F., Rivera, M., Roberts, L., Smallridge, R., Smyrk, T., Stanton, M., Thompson, R.H., Torbenson, M., Yang, J.D., Zhang, L., Brimo, F., Ajani, J.A., Gonzalez, A.M.A., Behrens, C., Bondaruk, J., Broaddus, R., Czerniak, B., Esmaeli, B., Fujimoto, J., Gershenwald, J., Guo, C., Lazar, A.J., Logothetis, C., Meric-Bernstam, F., Moran, C., Ramondetta, L., Rice, D., Sood, A., Tamboli, P., Thompson, T., Troncoso, P., Tsao, A., Wistuba, I., Carter, C., Haydu, L., Hersey, P., Jakrot, V., Kakavand, H., Kefford, R., Lee, K., Long, G., Mann, G., Quinn, M., Saw, R., Scolyer, R., Shannon, K., Spillane, A., Stretch, onathan, Synott, M., Thompson, J., Wilmott, J., Al-Ahmadie, H., Chan, T.A., Ghossein, R., Gopalan, A., Levine, D.A., Reuter, V., Singer, S., Singh, B., Tien, N.V., Broudy, T., Mirsaidi, C., Nair, P., Drwiega, P., Miller, J., Smith, J., Zaren, H., Park, J.-W., Hung, N.P., Kebebew, E., Linehan, W.M., Metwalli, A.R., Pacak, K., Pinto, P.A., Schiffman, M., Schmidt, L.S., Vocke, C.D., Wentzensen, N., Worrell, R., Yang, H., Moncrieff, M., Goparaju, C., Melamed, J., Pass, H., Botnariuc, N., Caraman, I., Cernat, M., Chemencedji, I., Clipca, A., Doruc, S., Gorincioi, G., Mura, S., Pirtac, M., Stancul, I., Tcaciuc, D., Albert, M., Alexopoulou, I., Arnaout, A., Bartlett, J., Engel, J., Gilbert, S., Parfitt, J., Sekhon, H., Thomas, G., Rassl, D.M., Rintoul, R.C., Bifulco, C., Tamakawa, R., Urba, W., Hayward, N., Timmers, H., Antenucci, A., Facciolo, F., Grazi, G., Marino, M., Merola, R., de Krijger, R., Gimenez-Roqueplo, A.-P., Piché, A., Chevalier, S., McKercher, G., Birsoy, K., Barnett, G., Brewer, C., Farver, C., Naska, T., Pennell, N.A., Raymond, D., Schilero, C., Smolenski, K., Williams, F., Morrison, C., Borgia, J.A., Liptay, M.J., Pool, M., Seder, C.W., Junker, K., Omberg, L., Dinkin, M., Manikhas, G., Alvaro, D., Bragazzi, M.C., Cardinale, V., Carpino, G., Gaudio, E., Chesla, D., Cottingham, S., Dubina, M., Moiseenko, F., Dhanasekaran, R., Becker, K.-F., Janssen, K.-P., Slotta-Huspenina, J., Abdel-Rahman, M.H., Aziz, D., Bell, S., Cebulla, C.M., Davis, A., Duell, R., Elder, J.B., Hilty, J., Kumar, B., Lang, J., Lehman, N.L., Mandt, R., Nguyen, P., Pilarski, R., Rai, K., Schoenfield, L., Senecal, K., Wakely, P., Hansen, P., Lechan, R., Powers, J., Tischler, A., Grizzle, W.E., Sexton, K.C., Kastl, A., Henderson, J., Porten, S., Waldmann, J., Fassnacht, M., Asa, S.L., Schadendorf, D., Couce, M., Graefen, M., Huland, H., Sauter, G., Schlomm, T., Simon, R., Tennstedt, P., Olabode, O., Nelson, M., Bathe, O., Carroll, P.R., Chan, J.M., Disaia, P., Glenn, P., Kelley, R.K., Landen, C.N., Phillips, J., Prados, M., Simko, J., Smith-McCune, K., VandenBerg, S., Roggin, K., Fehrenbach, A., Kendler, A., Sifri, S., Steele, R., Jimeno, A., Carey, F., Forgie, I., Mannelli, M., Carney, M., Hernandez, B., Campos, B., Herold-Mende, C., Jungk, C., Unterberg, A., von Deimling, A., Bossler, A., Galbraith, J., Jacobus, L., Knudson, M., Knutson, T., Ma, D., Milhem, M., Sigmund, R., Godwin, A.K., Madan, R., Rosenthal, H.G., Adebamowo, C., Adebamowo, S.N., Boussioutas, A., Beer, D., Giordano, T., Mes-Masson, A.-M., Saad, F., Bocklage, T., Landrum, L., Mannel, R., Moore, K., Moxley, K., Postier, R., Walker, J., Zuna, R., Feldman, M., Valdivieso, F., Dhir, R., Luketich, J., Pinero, E.M.M., Quintero-Aguilo, M., Carlotti, C.G., Dos Santos, J.S., Kemp, R., Sankarankuty, A., Tirapelli, D., Catto, J., Agnew, K., Swisher, E., Creaney, J., Robinson, B., Shelley, C.S., Godwin, E.M., Kendall, S., Shipman, C., Bradford, C., Carey, T., Haddad, A., Moyer, J., Peterson, L., Prince, M., Rozek, L., Wolf, G., Bowman, R., Fong, K.M., Yang, I., Korst, R., Rathmell, W.K., Fantacone-Campbell, J.L., Hooke, J.A., Kovatich, A.J., Shriver, C.D., DiPersio, J., Drake, B., Govindan, R., Heath, S., Ley, T., Van Tine, B., Westervelt, P., Rubin, M.A., Lee, J.I., Aredes, N.D., Mariamidze, A., 2018. The Immune Landscape of Cancer. Immunity 48, 812–830.e14. https://doi.org/10.1016/j.immuni.2018.03.023

25 Xiong, D., Wang, Y., You, M., 2020. A gene expression signature of TREM2hi macrophages and γδ T cells predicts immunotherapy response. Nat Commun 11, 5084. https://doi.org/10.1038/s41467-020-18546-x

26 Patz, F., n.d. Focal Pulmonary Evaluation with PET Scanning’ 5.

27 Warburg, O., 1956. On the Origin of Cancer Cells. Science, New Series 123, 309–314.

28 Chan, D.A., Sutphin, P.D., Nguyen, P., Turcotte, S., Lai, E.W., Banh, A., Reynolds, G.E., Chi, J.-T., Wu, J., Solow-Cordero, D.E., Bonnet, M., Flanagan, J.U., Bouley, D.M., Graves, E.E., Denny, W.A., Hay, M.P., Giaccia, A.J., n.d. Targeting GLUT1 and the Warburg Effect in Renal Cell Carcinoma by Chemical Synthetic Lethality 11.

29 Iansante, V., Choy, P.M., Fung, S.W., Liu, Y., Chai, J.-G., Dyson, J., Del Rio, A., D’Santos, C., Williams, R., Chokshi, S., Anders, R.A., Bubici, C., Papa, S., 2015. PARP14 promotes the Warburg effect in hepatocellular carcinoma by inhibiting JNK1-dependent PKM2 phosphorylation and activation. Nat Commun 6, 7882. https://doi.org/10.1038/ncomms8882

30 Jensen, D.H., Therkildsen, M.H., Dabelsteen, E., 2015. A reverse Warburg metabolism in oral squamous cell carcinoma is not dependent upon myofibroblasts. J Oral Pathol Med 44, 714–721. https://doi.org/10.1111/jop.12297

31 Witkiewicz, A.K., Whitaker-Menezes, D., Dasgupta, A., Philp, N.J., Lin, Z., Gandara, R., Sneddon, S., Martinez-Outschoorn, U.E., Sotgia, F., Lisanti, M.P., 2012. Using the “reverse Warburg effect” to identify high-risk breast cancer patients: Stromal MCT4 predicts poor clinical outcome in triple-negative breast cancers. Cell Cycle 11, 1108–1117. https://doi.org/10.4161/cc.11.6.19530

32 Whitaker-Menezes, D., Martinez-Outschoorn, U.E., Lin, Z., Ertel, A., Flomenberg, N., Witkiewicz, A.K., Birbe, R.C., Howell, A., Pavlides, S., Gandara, R., Pestell, R.G., Sotgia, F., Philp, N.J., Lisanti, M.P., n.d. Evidence for a stromal-epithelial “lactate shuttle” in human tumors. Cell Cycle 10, 12.

33 Najafi, M., Hashemi Goradel, N., Farhood, B., Salehi, E., Nashtaei, M.S., Khanlarkhani, N., Khezri, Z., Majidpoor, J., Abouzaripour, M., Habibi, M., Kashani, I.R., Mortezaee, K., 2019. Macrophage polarity in cancer: A review. J Cell Biochem 120, 2756–2765. https://doi.org/10.1002/jcb.27646

34 ChÃivez-GalÃin, L., Olleros, M.L., Vesin, D., Garcia, I., 2015. Much More than M1 and M2 Macrophages, There are also CD169+ and TCR+ Macrophages. Front. Immunol. 6. https://doi.org/10.3389/fimmu.2015.00263

35 Ferrer, G., Jung, B., Chiu, P.Y., Aslam, R., Palacios, F., Mazzarello, A.N., Vergani, S., Bagnara, D., Chen, S.-S., Yancopoulos, S., Xochelli, A., Yan, X.-J., Burger, J.A., Barrientos, J.C., Kolitz, J.E., Allen, S.L., Stamatopoulos, K., Rai, K.R., Sherry, B., Chiorazzi, N., 2021. Myeloid-derived suppressor cell subtypes differentially influence T-cell function, T-helper subset differentiation, and clinical course in CLL. Leukemia. https://doi.org/10.1038/s41375-021-01249-7

36 Kodach, L.L., Peppelenbosch, M.P., 2021. Targeting the Myeloid-Derived Suppressor Cell Compartment for Inducing Responsiveness to Immune Checkpoint Blockade Is Best Limited to Specific Subtypes of Gastric Cancers. Gastroenterology 161, 727. https://doi.org/10.1053/j.gastro.2021.03.047

37 Mu, X., Shi, W., Xu, Y., Xu, C., Zhao, T., Geng, B., Yang, J., Pan, J., Hu, S., Zhang, C., Zhang, J., Wang, C., Shen, J., Che, Y., Liu, Z., Lv, Y., Wen, H., You, Q., 2018. Tumor-derived lactate induces M2 macrophage polarization via the activation of the ERK/STAT3 signaling pathway in breast cancer. Cell Cycle 17, 428– 438. https://doi.org/10.1080/15384101.2018.1444305

38 Brand, A., Singer, K., Koehl, G.E., Kolitzus, M., Schoenhammer, G., Thiel, A., Matos, C., Bruss, C., Klobuch, S., Peter, K., Kastenberger, M., Bogdan, C., Schleicher, U., Mackensen, A., Ullrich, E., Fichtner-Feigl, S., Kesselring, R., Mack, M., Ritter, U., Schmid, M., Blank, C., Dettmer, K., Oefner, P.J., Hoffmann, P., Walenta, S., Geissler, E.K., Pouyssegur, J., Villunger, A., Steven, A., Seliger, B., Schreml, S., Haferkamp, S., Kohl, E., Karrer, S., Berneburg, M., Herr, W., Mueller-Klieser, W., Renner, K., Kreutz, M., 2016. LDHA-Associated Lactic Acid Production Blunts Tumor Immunosurveillance by T and NK Cells. Cell Metabolism 24, 657–671. https://doi.org/10.1016/j.cmet.2016.08.011

39 Cosmi, L., Annunziato, F., Galli, G., Manetti, R., Maggi, E., Romagnani, S., 2001. CRTH2: marker for the detection of human Th2 and Tc2 cells. Advances in experimental medicine and biology 495, 25.

40 Zheng, W., Flavell, R.A., 1997. The Transcription Factor GATA-3 Is Necessary and Sufficient for Th2 Cytokine Gene Expression in CD4 T Cells. Cell 89, 587–596. https://doi.org/10.1016/S0092-8674(00)80240-8

41 Mazzoni, A., Maggi, L., Siracusa, F., Ramazzotti, M., Rossi, M., Santarlasci, V., Montaini, G., Capone, M., Rossettini, B., De Palma, R., Kruglov, A., Cimaz, R., Maggi, E., Romagnani, S., Liotta, F., Cosmi, L., Annunziato, F., 2019. Eomes controls the development of Th17-derived (non-classic) Th1 cells during chronic inflammation. European Journal of Immunology 49, 79. https://doi.org/10.1002/eji.201847677

42 Gagliani, N., Vesely, M.C.A., Iseppon, A., Brockmann, L., Xu, H., Palm, N.W., de Zoete, M.R., Licona-Limón, P., Paiva, R.S., Ching, T., Weaver, C., Zi, X., Pan, X., Fan, R., Garmire, L.X., Cotton, M.J., Drier, Y., Bernstein, B., Geginat, J., Stockinger, B., Esplugues, E., Huber, S., Flavell, R.A., 2015. Th17 cells transdifferentiate into regulatory T cells during resolution of inflammation. Nature 523, 221–225. https://doi.org/10.1038/nature14452

43 Spranger, S., Bao, R., Gajewski, T.F., 2015. Melanoma-intrinsic β-catenin signalling prevents anti-tumour immunity. Nature 523, 231–235. https://doi.org/10.1038/nature14404

44 Tanaka, T., Kojima, K., Yokota, K., Tanaka, Y., Ooizumi, Y., Ishii, S., Nishizawa, N., Yokoi, K., Ushiku, H., Kikuchi, M., Kojo, K., Minatani, N., Katoh, H., Sato, T., Nakamura, T., Sawanobori, M., Watanabe, M., Yamashita, K., 2019. Comprehensive Genetic Search to Clarify the Molecular Mechanism of Drug Resistance Identifies ASCL2-LEF1/TSPAN8 Axis in Colorectal Cancer. Ann Surg Oncol 26, 1401–1411. https://doi.org/10.1245/s10434-019-07172-7

45 Sia, D., Jiao, Y., Martinez-Quetglas, I., Kuchuk, O., Villacorta-Martin, C., Castro de Moura, M., Putra, J., Camprecios, G., Bassaganyas, L., Akers, N., Losic, B., Waxman, S., Thung, S.N., Mazzaferro, V., Esteller, M., Friedman, S.L., Schwartz, M., Villanueva, A., Llovet, J.M., 2017. Identification of an Immune-specific Class of Hepatocellular Carcinoma, Based on Molecular Features. Gastroenterology 153, 812–826. https://doi.org/10.1053/j.gastro.2017.06.007

46 Chalmin, F., Mignot, G., Bruchard, M., Chevriaux, A., Végran, F., Hichami, A., Ladoire, S., Derangère, V., Vincent, J., Masson, D., Robson, S.C., Eberl, G., Pallandre, J.R., Borg, C., Ryffel, B., Apetoh, L., Rébé, C., Ghiringhelli, F., 2012. Stat3 and Gfi-1 Transcription Factors Control Th17 Cell Immunosuppressive Activity via the Regulation of Ectonucleotidase Expression. Immunity 36, 362–373. https://doi.org/10.1016/j.immuni.2011.12.019

47 Bengsch, B., Seigel, B., Flecken, T., Wolanski, J., Blum, H.E., Thimme, R., 2012. Human Th17 Cells Express High Levels of Enzymatically Active Dipeptidylpeptidase IV (CD26). J.I. 188, 5438–5447. https://doi.org/10.4049/jimmunol.1103801

48 Yang, X.O., Pappu, B.P., Nurieva, R., Akimzhanov, A., Kang, H.S., Chung, Y., Ma, L., Shah, B., Panopoulos, A.D., Schluns, K.S., Watowich, S.S., Tian, Q., Jetten, A.M., Dong, C., 2008. T Helper 17 Lineage Differentiation Is Programmed by Orphan Nuclear Receptors RORα and RORγ. Immunity 28, 29– 39. https://doi.org/10.1016/j.immuni.2007.11.016

49 49 Arra, A., Lingel, H., Kuropka, B., Pick, J., Schnoeder, T., Fischer, T., Freund, C., Pierau, M., Brunner-Weinzierl, M.C., 2017. The differentiation and plasticity of Tc17 cells are regulated by CTLA-4-mediated effects on STATs. OncoImmunology 6, e1273300. https://doi.org/10.1080/2162402X.2016.1273300

50 Chellappa, S., Hugenschmidt, H., Hagness, M., Subramani, S., Melum, E., Line, P.D., Labori, K.-J., Wiedswang, G., Taskén, K., Aandahl, E.M., 2017. CD8+ T Cells That Coexpress RORγt and T-bet Are Functionally Impaired and Expand in Patients with Distal Bile Duct Cancer. J. Immunol. 198, 1729. https://doi.org/10.4049/jimmunol.1600061

51 Heink, S., Yogev, N., Garbers, C., Herwerth, M., Aly, L., Gasperi, C., Husterer, V., Croxford, A.L., Möller-Hackbarth, K., Bartsch, H.S., Sotlar, K., Krebs, S., Regen, T., Blum, H., Hemmer, B., Misgeld, T., Wunderlich, T.F., Hidalgo, J., Oukka, M., Rose-John, S., Schmidt-Supprian, M., Waisman, A., Korn, T., 2017. Trans-presentation of IL-6 by dendritic cells is required for the priming of pathogenic TH17 cells. Nat Immunol 18, 74–85. https://doi.org/10.1038/ni.3632

52 Li, B., Jones, L.L., Geiger, T.L., 2018. IL-6 Promotes T Cell Proliferation and Expansion under Inflammatory Conditions in Association with Low-Level RORγt Expression. J. Immunol. 201, 2934. https://doi.org/10.4049/jimmunol.1800016

53 Fabre, T., Molina, M.F., Soucy, G., Goulet, J.-P., Willems, B., Villeneuve, J.-P., Bilodeau, M., Shoukry, N.H., 2018. Type 3 cytokines IL-17A and IL-22 drive TGF-β–dependent liver fibrosis. Sci. Immunol. 3, eaar7754. https://doi.org/10.1126/sciimmunol.aar7754

54 Kuang, D.-M., Peng, C., Zhao, Q., Wu, Y., Zhu, L.-Y., Wang, J., Yin, X.-Y., Li, L., Zheng, L., 2010. Tumor-Activated Monocytes Promote Expansion of IL-17–Producing CD8 + T Cells in Hepatocellular Carcinoma Patients. J.I. 185, 1544–1549. https://doi.org/10.4049/jimmunol.0904094

55 Kryczek, I., Bruce, A.T., Gudjonsson, J.E., Johnston, A., Aphale, A., Vatan, L., Szeliga, W., Wang, Y., Liu, Y., Welling, T.H., Elder, J.T., Zou, W., 2008. Induction of IL-17 + T Cell Trafficking and Development by IFN-γ: Mechanism and Pathological Relevance in Psoriasis. J Immunol 181, 4733–4741. https://doi.org/10.4049/jimmunol.181.7.4733

56 Patel, D.N., King, C.A., Bailey, S.R., Holt, J.W., Venkatachalam, K., Agrawal, A., Valente, A.J., Chandrasekar, B., 2007. Interleukin-17 Stimulates C-reactive Protein Expression in Hepatocytes and Smooth Muscle Cells via p38 MAPK and ERK1/2-dependent NF-κB and C/EBPβ Activation. J. Biol. Chem. 282, 27229–27238. https://doi.org/10.1074/jbc.M703250200

57 Sawa, S., Cherrier, M., Lochner, M., Satoh-Takayama, N., Fehling, H.J., Langa, F., Santo, J.P.D., Eberl, G., 2010. Lineage Relationship Analysis of RORgt+ Innate Lymphoid Cells 330, 6.

58 Zheng, Y., Valdez, P.A., Danilenko, D.M., Hu, Y., Sa, S.M., Gong, Q., Abbas, A.R., Modrusan, Z., Ghilardi, N., de Sauvage, F.J., Ouyang, W., 2008. Interleukin-22 mediates early host defense against attaching and effacing bacterial pathogens. Nat Med 14, 282–289. https://doi.org/10.1038/nm1720

59 Newman, A.M., Steen, C.B., Liu, C.L., Gentles, A.J., Chaudhuri, A.A., Scherer, F., Khodadoust, M.S., Esfahani, M.S., Luca, B.A., Steiner, D., Diehn, M., Alizadeh, A.A., 2019. Determining cell type abundance and expression from bulk tissues with digital cytometry. Nat Biotechnol 37, 773–782. https://doi.org/10.1038/s41587-019-0114-2

60 https://www.ncbi.nlm.nih.gov/gap

61 Chen, P.-L., Roh, W., Reuben, A., Cooper, Z.A., Spencer, C.N., Prieto, P.A., Miller, J.P., Bassett, R.L., Gopalakrishnan, V., Wani, K., De Macedo, M.P., Austin-Breneman, J.L., Jiang, H., Chang, Q., Reddy, S.M., Chen, W.-S., Tetzlaff, M.T., Broaddus, R.J., Davies, M.A., Gershenwald, J.E., Haydu, L., Lazar, A.J., Patel, S.P., Hwu, P., Hwu, W.-J., Diab, A., Glitza, I.C., Woodman, S.E., Vence, L.M., Wistuba, I.I., Amaria, R.N., Kwong, L.N., Prieto, V., Davis, R.E., Ma, W., Overwijk, W.W., Sharpe, A.H., Hu, J., Futreal, P.A., Blando, J., Sharma, P., Allison, J.P., Chin, L., Wargo, J.A., 2016. Analysis of Immune Signatures in Longitudinal Tumor Samples Yields Insight into Biomarkers of Response and Mechanisms of Resistance to Immune Checkpoint Blockade. Cancer Discov 6, 827. https://doi.org/10.1158/2159-8290.CD-15-1545

62 Bonuccelli, G., Whitaker-Menezes, D., Castello-Cros, R., Pavlides, S., Pestell, R.G., Fatatis, A., Witkiewicz, A.K., Vander Heiden, M.G., Migneco, G., Chiavarina, B., Frank, P.G., Capozza, F., Flomenberg, N., Martinez-Outschoorn, U.E., Sotgia, F., Lisanti, M.P., 2010. The reverse Warburg Effect: Glycolysis inhibitors prevent the tumor promoting effects of caveolin-1 deficient cancer associated fibroblasts. Cell Cycle 9, 1960–1971. https://doi.org/10.4161/cc.9.10.11601

63 Saha, D., n.d. Macrophage Polarization Contributes to Glioblastoma Eradication by Combination Immunovirotherapy and Immune Checkpoint Blockade 21.

64 Durbin, B.P., Hardin, J.S., Hawkins, D.M., Rocke, D.M., 2002. A variance-stabilizing transformation for gene-expression microarray data. Bioinformatics 18, S105–S110. https://doi.org/10.1093/bioinformatics/18.suppl_1.S105

65 Li, X., Brock, G.N., Rouchka, E.C., Cooper, N.G.F., Wu, D., O’Toole, T.E., Gill, R.S., Eteleeb, A.M., O’Brien, L., Rai, S.N., 2017. A comparison of per sample global scaling and per gene normalization methods for differential expression analysis of RNA-seq data. PLoS ONE 12, e0176185. https://doi.org/10.1371/journal.pone.0176185

